# The Dynamic Strength of the Hair-Cell Tip Link Reveals Mechanisms of Hearing and Deafness

**DOI:** 10.1101/763847

**Authors:** Eric M. Mulhall, Andrew Ward, Darren Yang, Mounir A. Koussa, David P. Corey, Wesley P. Wong

## Abstract

Our senses of hearing and balance rely on the extraordinarily sensitive molecular machinery of the inner ear to convert deflections as small as the width of a single carbon atom^1,2^ into electrical signals that the brain can process^3^. In humans and other vertebrates, transduction is mediated by hair cells^4^, where tension on tip links conveys force to mechanosensitive ion channels^5^. Each tip link comprises two helical filaments of atypical cadherins bound at their N-termini through two unique adhesion bonds^6–8^. Tip links must be strong enough to maintain a connection to the mechanotransduction channel under the dynamic forces exerted by sound or head movement—yet might also act as mechanical circuit breakers, releasing under extreme conditions to preserve the delicate structures within the hair cell. Previous studies have argued that this connection is exceptionally static, disrupted only by harsh chemical conditions or loud sound^9–12^. However, no direct mechanical measurements of the full tip-link connection have been performed. Here we describe the dynamics of the tip-link connection at single-molecule resolution and show how avidity conferred by its double stranded architecture enhances mechanical strength and lifetime, yet still enables it to act as a dynamic mechanical circuit breaker. We also show how the dynamic strength of the connection is facilitated by strong cis-dimerization and tuned by extracellular Ca^2+^, and we describe the unexpected etiology of a hereditary human deafness mutation. Remarkably, the connection is several thousand times more dynamic than previously thought, challenging current assumptions about tip-link stability and turnover rate, and providing insight into how the mechanotransduction apparatus conveys mechanical information. Our results reveal fundamental mechanisms that underlie mechanoelectric transduction in the inner ear, and provide a foundation for studying multi-component linkages in other biological systems.

## Main Text

The tip link is a double-stranded complex of the Ca^2+^-dependent extracellular adhesion proteins protocadherin-15 (PCDH15) and cadherin-23 (CDH23), bound at their N-termini (Fig. 1a)^6,8^. While classic *trans* cadherin bonds are mediated by homophilic strand swapping of an N-terminal tryptophan residue into a hydrophobic pocket of its binding partner^13^, we previously demonstrated that the *trans* tip-link bond has a far more extensive, amphiphilic “handshake” interface involving ~30 amino acids in the first two extracellular (EC) domains of each protein^8^. Although the dissociation constant of a single tip-link bond is approximately the same as for classic cadherins^8,14–18^, little is known about how the tip-link bond responds to force.

**Fig. 1:**
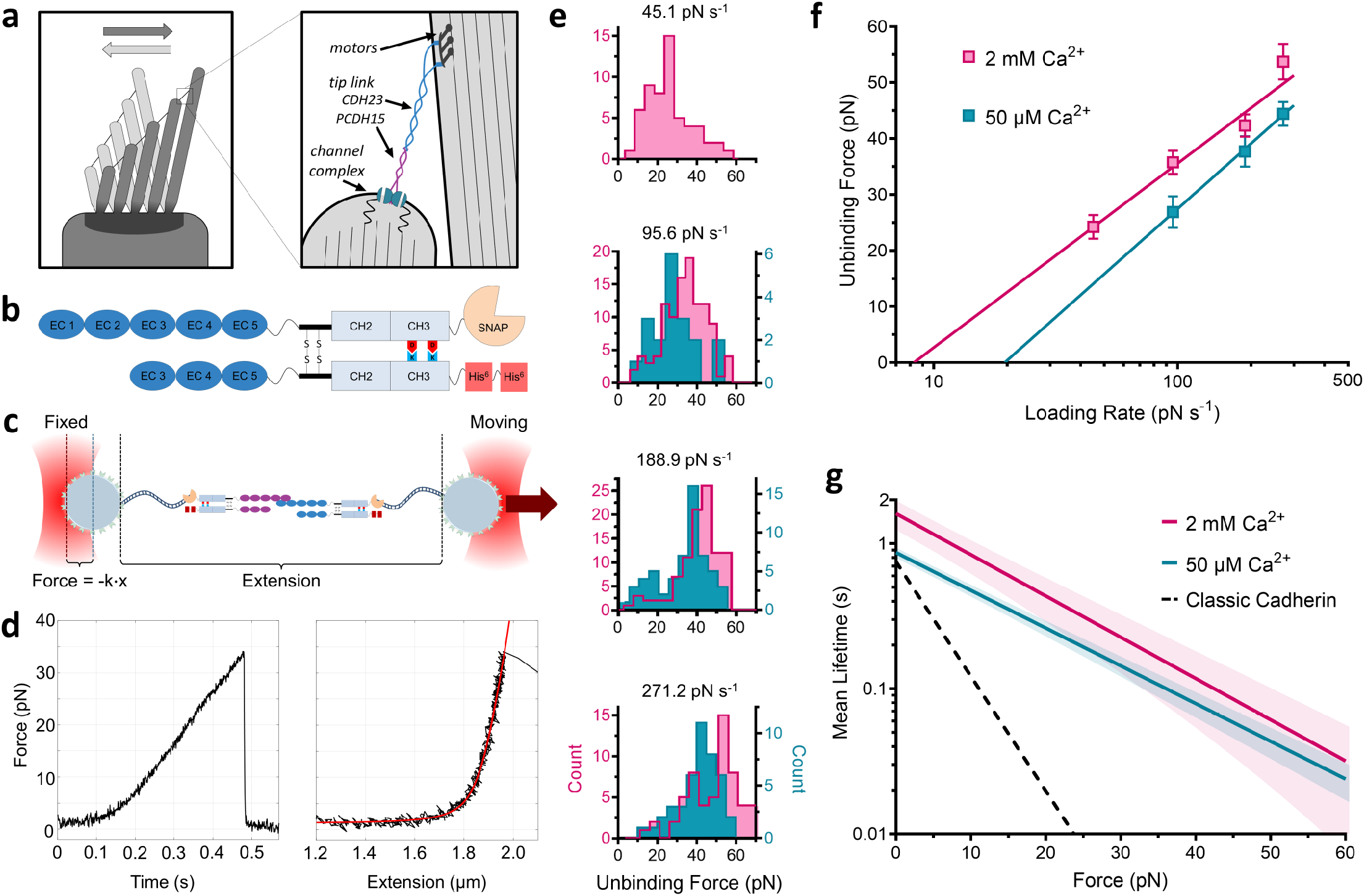
Resistance of the tip-link bond to mechanical stress and Ca^2+^-poor cochlear endolymph. **a**, The hair-cell stereocilia bundle and mechanotransduction complex. The mechanotransduction complex is shown with a single dimeric tip-link filament connecting to a dimeric transduction channel^56^. **b**, Single-bond fusion proteins containing one EC1-2 binding domain. **c**, Single-bond fusion proteins in the dual-beam optical trap. Force was calculated as a linear function of displacement from a stiffness-calibrated optical trap. Extension was measured as the distance between the two surfaces of each microsphere. **d**, Representative force-time and force-extension profiles for a single-bond unbinding event. The force-time profile was used to extract the force loading rate in the linear regime between 0.2 s and the unbinding event at 0.47 s. The force-extension profile was fit with an extensible worm-like chain (WLC) model to extract contour length, persistence length, and unbinding force for each unbinding trace. **e**, Histograms of unbinding forces at different force loading rates in 50 μM and 2 mM Ca^2+^. **f**, Most-probable unbinding forces plotted as a function of loading rate. A kernel density estimation was used to determine the most likely unbinding force for each condition. Error shown as the optimal kernel bandwidth. A linear fit of the data was used to extract force-dependent kinetics^22^. **g**, Mean single-bond lifetime as a function of force calculated using *k_off_^0^* and *f_β_* from **f**, compared to the average lifetime of classic cadherin bonds (dashed black line)^14–18^. The propagated errors in lifetime due to errors in fit parameters from **f** are shown as light bands.

The tip link’s double-stranded architecture is also distinct from classic cadherins, which form clusters of tens or hundreds of monomeric proteins in a variety of tissues^19^ and rely on the additive strength of many links to maintain their connection over hours, days, or weeks^20,21^. Tip links must be strong enough to maintain the connection to the mechanotransduction channel under the dynamic forces exerted by sound or head movement, yet they also disassemble during exposure to loud noise, ostensibly to protect the mechanotransduction apparatus or the integrity of the post-mitotic hair cell^9^. How can a linkage with only two strands faithfully perform these tasks? Although studies exploring the recovery time of tip links in response to disruption via Ca^2+^ chelation or noise damage have proposed that the lifetime of the tip-link connection may be on the order of hours^9–12^, no biophysical measurements have established the strength or dynamics of this linkage.

To measure the mechanical strength of an individual tip-link bond, we used optical tweezers to perform single-molecule dynamic force spectroscopy on engineered fusion proteins (Fig. 1b-d) (Extended Data Fig. 1–5). Unbinding force data were fit with the Bell-Evans model^22^ to reveal several salient features (Fig. 1e-f). The mean lifetime (1/*k_off_*) of the single tip-link bond at zero force is 1.6 ± 0.4 seconds (2 mM Ca^2+^, *k_off_^0^* = 0.62 ± 0.21 s^−1^) (Fig. 1g), agreeing well with independent zero-force measurements^8,22,23^. Force generally destabilizes adhesion bonds by lowering the energy barrier for unbinding, decreasing the time that two proteins will remain bound^24^. The force sensitivity of the tip-link bond is *f_β_* = 14.3 ± 5.0 pN, the characteristic force required to accelerate the off-rate by *e*-fold (Fig. 1f). Compared with classic cadherins (mean lifetime ≈ 0.75 s, *f_β_* ≈ 5.5 pN)^14–18^, the single tip-link bond is significantly more resistant to mechanical force (Fig. 1g).

The tip-link bond is also destabilized by very low [Ca^2+^]^8,25^. However, we found that the mean lifetime of a single tip-link bond is reduced by only 46% in 50 μM Ca^2+^ (mean zero-force lifetime = 0.87 ± 0.07 s), a concentration near that of the endolymph fluid that bathes tip links in the cochlea (Fig. 1g). This insensitivity to low [Ca^2+^] is in stark contrast with classic cadherins, for which half-maximal adhesion of even the relatively Ca^2+^-insensitive N-cadherin is observed at 720 μM Ca^2+^ ^26^. Thus, the tip-link bond has evolved independently of classic cadherin bonds to be both more resistant to the force experienced during mechanotransduction and less sensitive to the Ca^2+^-poor endolymph, properties which enable the tip link to effectively convey mechanical stimuli in the inner ear.

Why is the tip link double stranded? Avidity is the cumulative strength that results from multiple bonds, and one important contribution is the division of externally applied force among the individual components^27^. Force-sharing among bonds can result in dramatic increases in lifetime because the off-rate of a single bond has a roughly exponential dependence on force. Another contribution comes from the ability of individual components to rebind after they rupture, dynamically maintaining the overall connection. Since CDH23 and PCDH15 both form strong cis-dimers *in-vitro* and *in-vivo*, facilitated by bonds along their ectodomains^6,7,28–31^, the tip link may increase its lifetime through rebinding if opposing PCDH15 and CDH23 EC 1-2 domains are kept in sufficiently close spatial proximity, creating a high effective local concentration and a fast on-rate.

The tip-link connection may be described by a kinetic model (Fig. 2a) (Supplementary Discussion) in which a transition from the double-bound (B2) to unbound (U) state requires an intermediate single-bound (B1) state. If opposing PCDH15 and CDH23 EC1-2 domains are spatially constrained by cis-dimerization interfaces^28^, the effective concentration-dependent binding rate *C_eff_k_on_* of an unbound pair might be sufficiently fast relative to the single bond off-rate *k_off_* to allow the bond to transition back to the B2 state before moving to the U state where complete rupture occurs. The effective local concentration *C_eff_* can be approximated by calculating the volume of a sphere with a radius equal to the length an individual EC1-2 binding domain (~10 nm)^8^, yielding *C_eff_* ~400 μM (Supplementary Discussion). Using a single-bond *k_on_* of 6.2 ± 2.8 x 10^4^ M^−1^ s^−1^ ^32^ and a single-bond *k_off_* of 0.6 s^−1^ (Fig. 1g), we estimate the zero-force rebinding rate (B1➔B2) to be ~25 ± 14 s^−1^, and the overall lifetime of the connection to be 17-52 seconds, much longer than the single-bond lifetime calculated above of 1.6 ± 0.4 seconds.

**Fig. 2:**
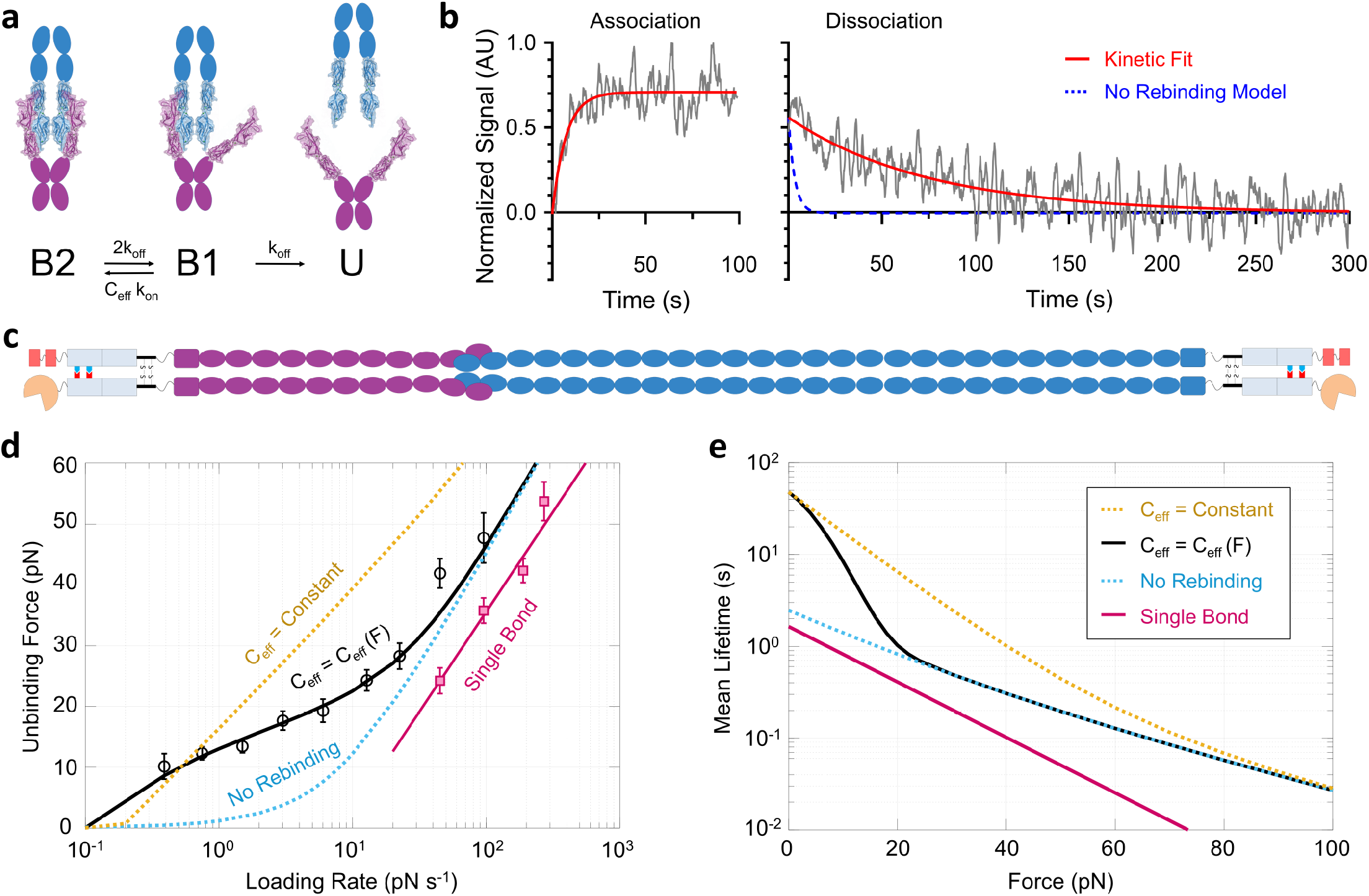
Dynamics of the double-stranded tip-link bond. **a**, State diagram for tip-link avidity. **b**, Biolayer interferometry of dimeric full-ectodomain PCDH15 and CDH23 fusion proteins. *left*, Association enables calculation of *C_eff_k_on_. right*, Dissociation rate fit (red) indicates a lifetime much longer than that calculated without re-binding (blue dashed line). Here, 320 nM of both PCDH15 and EC1-2 truncated PCDH15 were used for the association phase. **c**, Schematic of dimeric, full-ectodomain proteins used for biolayer interferometry and force-spectroscopy experiments. PCDH15 is shown in purple and CDH23 in blue; Fc domains, SNAP tag and tandem His6 tag as in Fig. 1b. **d**, Binding avidity of full-length dimers revealed at low force. Most-probable unbinding forces for double-bond full-length dimer proteins in 2 mM Ca^2+^ (black circles); single-bond unbinding forces from Fig. 1f (magenta squares). A best-fit kinetic model to both the single-bond and double-bond interactions is shown as solid lines. A no-rebinding model is shown as the blue dashed line; a constant-*C_eff_* model (fixed at the zero-force value of 800 μM determined from biolayer interferometry) is shown as the orange dashed line. **e**, Calculated mean lifetime from the model fits in **d**. At low forces, the tip link has a pronounced increase in lifetime relative to the single bond due to avidity. At high forces, resistance to force is only doubled relative to that of a single bond, due to force sharing between the two bonds.

To test this hypothesis, we used biolayer interferometry^33,34^ to measure the zero-force kinetics of the double-bond tip link interaction (Fig. 2c, Extended Data Fig. 6). We found the mean lifetime of the interaction (1/*k_off_*) to be 71.1 ± 5.3 s (SEM, *n* = 18) and the on-rate to be 1.32 ± 0.5 x 10^5^ M^−1^ s^−1^ (Fig. 2b) in reasonable agreement with our estimates and with published single-bond on-rates^32^. These values provide an empirical means of determining the effective concentration *C_eff_* in the B1 state, yielding *C_eff_* ~800 ± 400 μM, three orders of magnitude more concentrated than the equilibrium K_D_ of a single-bond interaction^8,23^ and sufficient for promoting rapid rebinding to the B2 state (Supplementary Discussion). Thus, at zero force the tip-link lifetime is substantially increased by rebinding, facilitated by cis-dimerization domains which keep binding domains in close proximity and the effective concentration high.

In the inner ear, the tip link is subject to a resting tension of ~10 pN applied through myosin motors attached intracellularly to CDH23^35,36^, and may experience forces exceeding 100 pN from high intensity sound^37^. To determine the dynamics of the tip-link connection under force, we used dynamic force spectroscopy to measure the unbinding kinetics of the double-stranded tip link containing the entire ectodomains (Fig. 2c) (Extended Data Table 2). We found a large increase in strength (Fig. 2d, black circles) relative to the single bond (magenta squares). From single-bond data (Fig. 1f), we can independently predict the rupture force in the case of simple load-sharing between two bonds^27^ (Fig. 2d, “*No Rebinding*”) (Supplementary Discussion). Remarkably, in the force regime of 10-25 pN, in which the tip link normally operates^35,38^, measured rupture forces are significantly higher than predicted by simple load sharing, suggesting that rebinding occurs even under force.

A rebinding model, based on single-bond kinetics and a *C_eff_* that was determined from biolayer interferometry and is assumed to be independent of force, successfully predicts higher unbinding forces, but it predicts forces even higher than observed (Fig. 2d, “*C_eff_ = Constant*”) (Supplementary Discussion). Why? Recent studies using optical tweezers and molecular dynamics simulations suggest that tip link proteins are far more elastic than initially thought^39,40^. Elasticity will allow unbound domains in the B1 state (Fig. 2a) to separate as force increases, decreasing *C_eff_* and reducing rebinding. A three-state kinetic model in which *C_eff_* decreases with a Gaussian decay dependent on the compliance parameter *f_c_* (Supplementary Discussion)^41^ fits the data well (Fig. 2d, *“C_eff_ = C_eff_*(*F*)”), yielding *f_c_* = 10.4 ± 2.0 pN and an effective concentration at zero force of *C_eff_^0^* = 479 ± 200 μM. This model predicts a zero-force mean lifetime of 55 ± 43 s, in good agreement with both our biolayer interferometry measurements, and our estimates based on extrapolating the effect of avidity from single-bond measurements. The model also predicts that elasticity is an important factor is determining the dynamics of the tip-link connection within the physiological range of forces experienced by the mechanotransduction apparatus.

Importantly, these measurements enable the prediction of mean tip-link lifetimes as a function of force (Fig. 2e, Extended Data Fig. 7) for different force histories. At a constant resting tension of 10 pN^30^, the double-bond interaction lasts an order of magnitude longer than a single-bond interaction (black line). Yet at very high forces of 50-60 pN the predicted lifetime is much less than a second. The tip link has apparently evolved mechanisms to stabilize the connection at the normally low operating forces, but to release quickly at higher, potentially damaging forces in order to protect the mechanotransduction complex and to reduce the magnitude of transduction.

Since our model predicts that the mechanical properties of tip-link cadherins significantly affect avidity, we made structural modifications to the tip link proteins to test this effect. Cis-dimerization links PCDH15 and CDH23 laterally at several points along their length^6,7,28–31^, and may act to stabilize *C_eff_* under force. We truncated PCDH15 and CDH23 to their first five EC domains in an effort to disrupt full cis-dimerization, and used single-molecule force spectroscopy to determine the effects on bond strength (Fig. 3a). At higher loading rates (> 20 pN/s), EC1-5 dimers displayed an enhanced strength relative to the single-bond interaction, very similar to the full dimer constructs. However, at slower loading rates the truncated dimer ruptured at lower forces than the full dimer, exhibiting a lifetime similar to that predicted by a simple force-sharing dimer model (blue line). This suggests that for the truncated constructs, rebinding is not occurring at forces above 6 pN (Extended Data Table 1–2). These data highlight the importance of cis-dimerization in enhancing the strength of the tip-link connection.

**Fig. 3.**
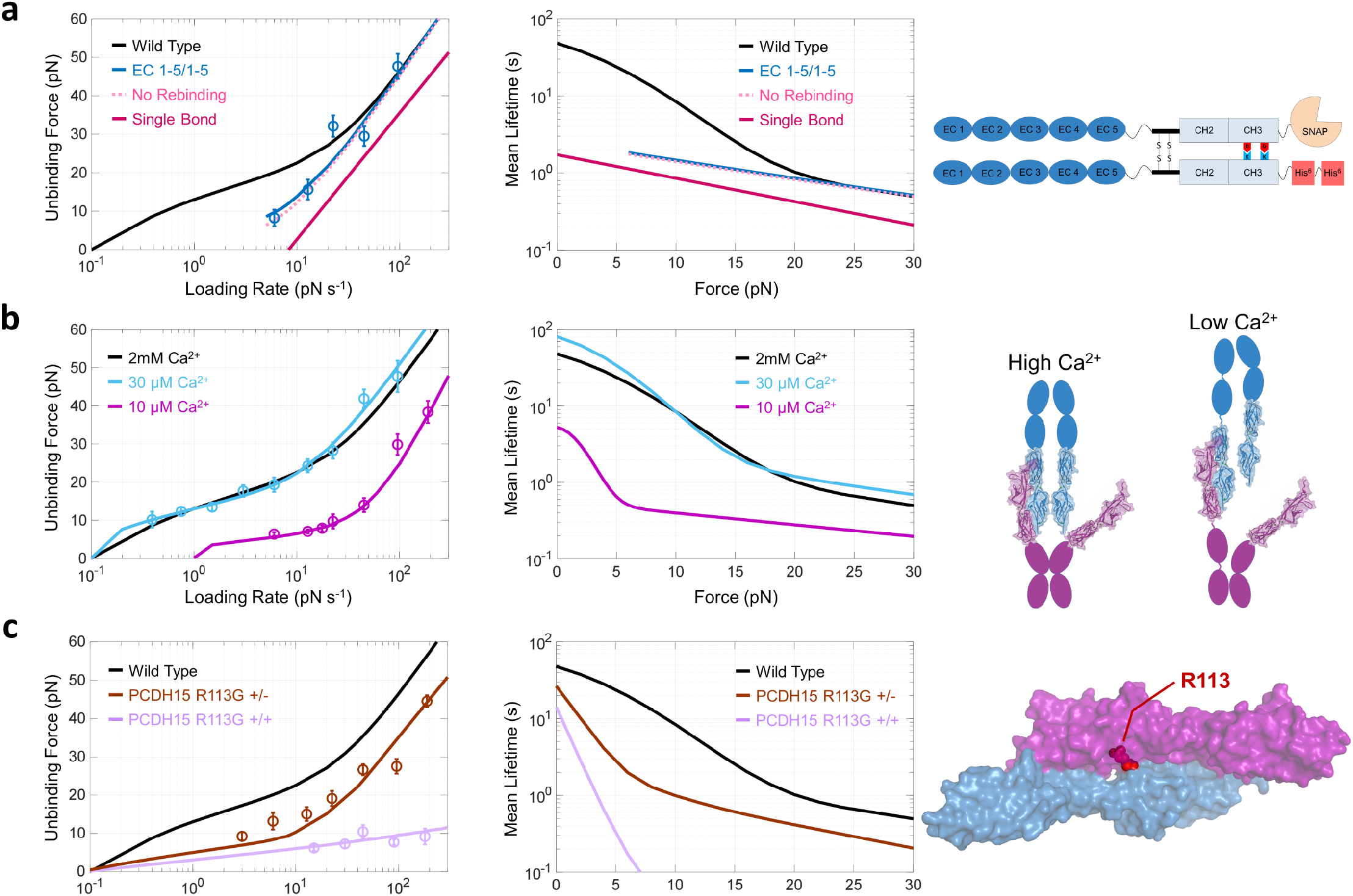
Tip-link dynamics modified by cis-dimerization interfaces, [Ca^2+^] and deafness mutations. **a**, Avidity under force is enhanced by cis-dimerization. *right*, Dimeric tip-link proteins were truncated to just the first five EC domains, to delete cis-dimerization interfaces further from the N-termini. Rupture data (blue circles) were fit with the force-dependent avidity model, with fixed parameters *k_off_* = 0.62 s^−1^ and *f_β_* = 14.3 pN (EC1-5/1-5). At the rupture forces tested, the model fit was indistinguishable from a model in which there is no rebinding. Truncated proteins only show an enhancement in strength attributable to force sharing. **b**, Low [Ca^2+^] destabilizes the tip link both by accelerating the single-bond off-rate (10 μM Ca^2+^, *k_off_* = 2.6 ± 0.8 s^−1^) and by altering the mechanical properties of the protein complex (10 μM Ca^2+^, *f_c_* = 2.9 ± 1.1 pN). *right*, A schematic of the tip link illustrating the consequence of increased protein compliance on *C_eff_*. **c**, The human deafness mutation R113G in PCDH15 (*right*; PDB 4AXW) disrupts the tip-link connection by altering force sensitivity. Rupture data from dimeric, full-ectodomain proteins, with one (brown) or both (purple) PCDH15 strands mutated, were simultaneously fit with a four-state model (Extended Data Figure 6a, Supplementary Discussion). The R113G mutation accelerates the zero-force off-rate by a factor of 2.3 and also makes the off-rate ~8 times more sensitive to force.

Ca^2+^ ions link the EC domains in each tip-link protein^8,42^, and act to maintain its structural integrity^39^. Chelation of extracellular Ca^2+^ disrupts tip links in intact hair cells^25^, and destabilizes the CDH23-PCDH15 interaction^8^ (Fig. 1g) (Extended Data Fig. 6c). As the concentration of Ca^2+^ bathing the tip link decreases, the junctions between each EC domain become more flexible due to decreased occupancy of Ca^2+^ ^8^, and the series elasticity of EC domains is increased^40^. We showed that protein elasticity modifies the extent of avidity under force (Fig. 2d). Therefore, increased compliance in low Ca^2+^ should cause unbound EC domains in the B1 state (Fig. 2a) to separate more rapidly as force increases, decreasing *C_eff_* and reducing rebinding.

In 30 μM Ca^2+^, the concentration found in bulk cochlear endolymph^43,44^, the lifetime of the tip-link connection at all forces was only moderately decreased relative to 2 mM Ca^2+^ (Fig. 3b, Extended Data Fig. 6c), again demonstrating that the tip link is well equipped to convey mechanical stimuli at physiologic concentrations of Ca^2+^. However, at the sub-endolymphatic level of 10 μM Ca^2+^, the tip link bond was severely weakened under force (Fig. 3b). Based on model fits to our force spectroscopy data, the single-bond off-rate at this calcium concentration is accelerated and the protein compliance is increased ~10 fold (Extended Data Table 1–2), causing *C_eff_* to decrease rapidly with force. Since the concentration of Ca^2+^ bathing the tip link is highly variable, and changes dynamically with magnitude of stimulation, for instance from loud sounds^45,46^, these cellular and biophysical properties provide an opportunity for cellular regulation of tip link resilience—and transduction magnitude—through extracellular [Ca^2+^].

A broad spectrum of Usher syndrome mutations disrupt tip-link integrity in hair cells^47,48^. The human PCDH15 R113G mutation causes hearing loss without a vestibular or visual phenotype^49^. The R113 residue participates in the binding interface, and the R113G mutation reduces bond affinity *in vitro^8,32^*. To determine how this mutation disrupts function, we used polarized Fc-domain dimerization to introduce this mutation either heterozygously (R113G^+/-^) or homozygously (R113G^+/+^), and measured its strength with force spectroscopy (Fig. 3c). The heterozygous R113G^+/-^ tip-link bond (brown circles) is weaker than the wild-type, and the homozygous R113G^+/+^ bond (purple circles) is much weaker at all forces (Extended Data Fig. 6). The exceptional reduction in strength by a single amino acid substitution is additional confirmation that our measurements are specific to the tip-link bond.

To understand how this mutation disrupts function, we performed a simultaneous fit of R113G^+/-^ and R113G^+/+^ unbinding forces, using a modified four-state force-dependent model (Extended Data Fig. 8) (Supplementary Discussion). Most strikingly, the R113G mutation causes the single-bond off-rate to be ~8 times more sensitive to force (f_β_^R113G^ = 1.8 ± 0.3 pN) compared with the wild-type bond (f_β_^WT^ = 14.3 ± 5.0 pN), without altering the force-dependence of *C_eff_* (Extended Data Table 1–2, Fig. 8c). Our results indicate the R113G^+/+^ protein remains bound to CDH23 11% as long as the wild type at zero force, but lasts just 0.3% as long at a resting tension of 10 pN. Since cochlear hair cells experience considerably more force than vestibular hair cells or the calyceal processes of retinal photoreceptors^50^, PCDH15 R113G^+/+^ likely unbinds more rapidly in the cochlea, explaining the more profound deafness phenotype in PCDH15 R113G^+/+^ human patients compared with the balance or visual phenotype.

*In vivo*, auditory stimuli oscillate hair bundles with a sinusoidal waveform^51^, so we performed Monte Carlo simulations to determine how oscillatory stimuli affect mean lifetime of the normal double-bond tip-link (Fig. 4). Using our empirically-determined parameters for the force-dependent kinetics of tip link bonds, we find that the mean lifetime of the connection is essentially insensitive to changes in sound frequency within the normal operating range (Fig. 4d) (~0-20 pN, ~20 Hz – 10 kHz) of the transduction complex. This effect is largely facilitated by efficient re-binding from state B1➔B2 during the slack half-cycle of the sine wave (Supplemental Movie 1). At very large stimulus magnitudes, corresponding to damaging noise stimuli^52^, the lifetime is significantly reduced. When we further include “slow” adaptation, mediated by the slipping/climbing of myosin motors that act to maintain resting tension^53,54^, tip-link lifetime is significantly enhanced by the relaxation of the peak forces and reduction of average force (Fig. 4e) (Supplemental Discussion). Overall, these surprising results indicate that multiple effects act to maintain an essentially constant lifetime of the connection at physiologically relevant frequencies and amplitudes.

**Fig. 4.**
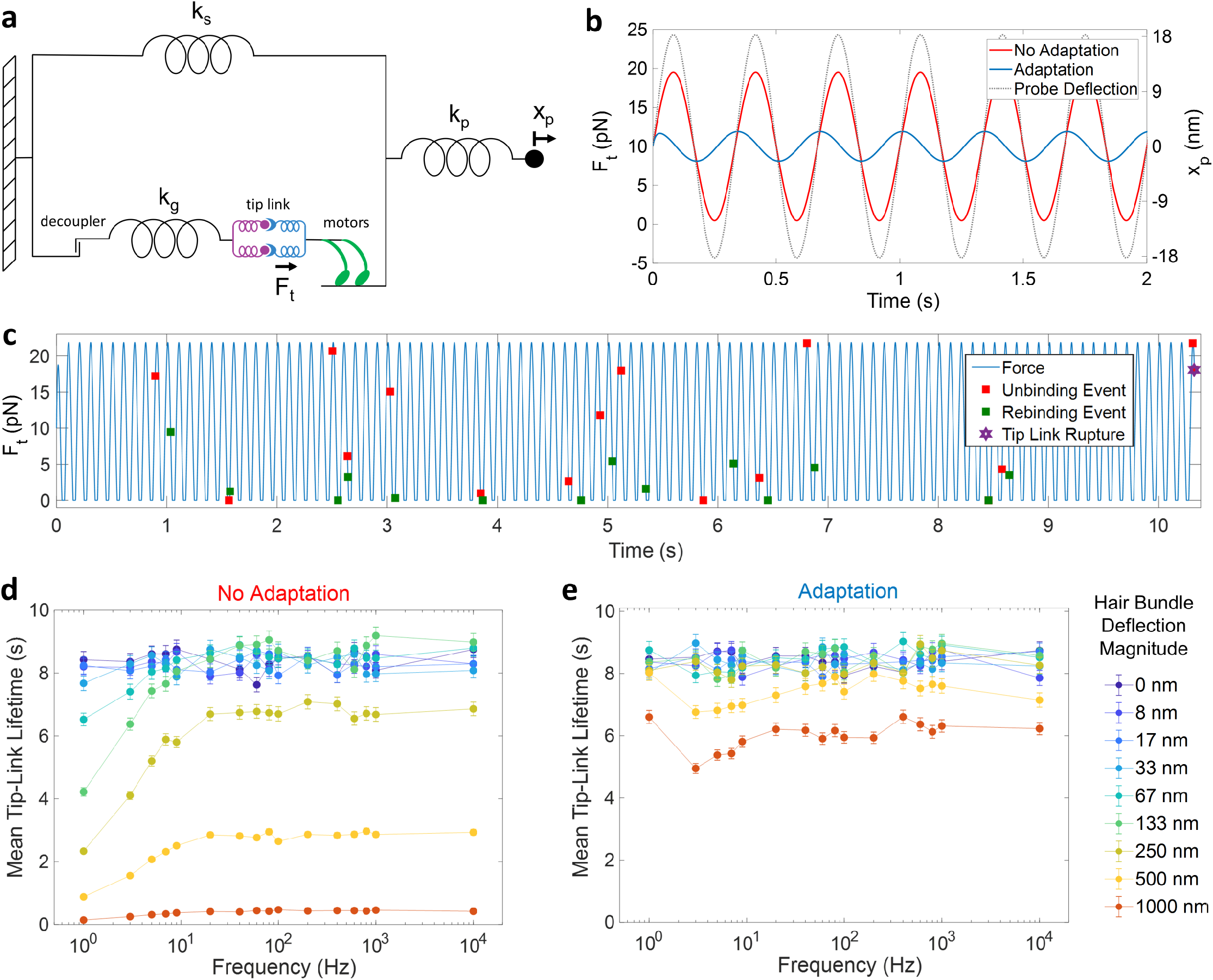
The lifetime of the tip-link connection in response to physiologically relevant oscillatory strain. **a**, A schematic of the modified mechanical model^53^ (Supplementary Discussion) used to calculate mean tip-link lifetime. k_s_ is the pivot stiffness of the stereocilia bundle, k_g_ is the gating spring stiffness, k_p_ is the probe stiffness, x_p_ is the probe deflection magnitude, and F_t_ is the force on the tip link. Hair bundle deflection magnitude is xp divided by γ = 0.12^38,54^. **b**, Force on an individual tip link F_t_ as a function of time at a 3 Hz stimulus frequency with and without “slow” adaptation (x_p_ = 18 nm). **c**, A single Monte Carlo simulation trajectory of tip-link lifetime performed using force-dependent off-rates and concentration-dependent on-rates obtained from force spectroscopy of full-length dimers (Fig. 2d-e). Unbinding and rebinding events are marked with squares. _d_, Mean tip-link lifetimes, based on thousands of simulations at frequencies between 1-10,000 Hz and at hair bundle deflection magnitudes of 0-1000 nm (error bars = SEM, *n* = 5,000). Without adaptation, tip-link lifetime is insensitive to both frequency and amplitude within the physiological range of hearing in humans (~20-10,000 Hz) for hair bundle deflections up to 133 nm, corresponding to a peak force of ~9 pN above resting tension on the tip link. **e**, Adaptation increases mean tip-link lifetime by reducing the force amplitude of each oscillation via slippage of the adaptation motors (Supplementary Discussion).

Although previous studies have depicted the tip link as a relatively static structure^11,12,55^, our results indicate the tip link is a highly dynamic connection. We estimate that tip-link lifetimes under physiological oscillatory forces (Fig. 4, Supplementary Discussion) are nearly 3000 times shorter than predicted from recovery experiments. We propose that the fast on-rate and close proximity of unbound cadherins could allow the tip link to re-form quickly across the gap between stereocilia, perhaps in tenths of a second (Extended Data Fig. 9), (Supplementary Discussion). Under normal conditions, we expect the majority of tip links to be bound and to be available to transduce force. During exposure to loud sounds, high forces likely shift the equilibrium distribution of tip-link connections towards the unbound state, reducing transduction amplitude, and acting as a type of dynamic mechanical circuit breaker.

The inner ear serves as a gateway for mechanical information from our environment. Our perception of sound and head movement is mediated by mechanotransduction complexes that convert nanometer-scale deflections into electrical signals. Yet, how this apparatus functions at the level of individual molecules has been poorly understood. Here, we studied the biophysical dynamics of the tip-link connection with single-molecule resolution and found that it has multiple mechanisms—distinct from classic cadherin linkages—that facilitate mechanotransduction in the inner ear. These mechanisms include a large bond interface to enhance strength, a double-stranded architecture mediated by strong cis-dimerization that permits rebinding before rupture, and tunable force- and Ca^2+^-dependent elastic properties that modify the lifetime of the connection. Strikingly, we also find this connection is over four orders of magnitude more dynamic than previously described, challenging the current view of the tip link as a static structure. For tip links, avidity is key to their strength, and our work reveals how Nature uses force distributed across multiple bonds to create a tunable mechanical circuit breaker—a motif that is likely used in other natural systems, and could be used as a design principle in synthetic systems as well.

## Methods

### Protein production and purification

Using antibody Fc heterodimers^57,58^, we engineered dimeric mouse PCDH15 and CDH23 proteins containing their first five EC domains, and we restricted binding to a single PCDH15-CDH23 bond by removing EC1-2 from one protein of each parallel dimer (Fig. 1b). This design was essential: monomeric EC1-2 and EC1-5 fusion proteins without the paired EC3-5 were unstable and bound non-specifically^8^. For single-bond tip link constructs with one EC1-2 binding domain, the coding regions of EC1-5 for mouse Pcdh15 and Cdh23 were amplified by PCR from full-length cDNA clones. This corresponds to base pairs 422-2269 from Pcdh15 RefSeq cDNA NM_023115 and base pairs 408-2066 from Cdh23 RefSeq cDNA NM_023370. Fusion proteins containing only the EC1-2 binding domains for each protein were unstable and bound non-specifically, and the addition of EC3-5 was necessary to stabilize these proteins, as determined from both biolayer interferometry and force spectroscopy. The EC3-5 domains were created by deleting amino acids 28-262 for PCDH15 and 25-228 for CDH23 using site-directed mutagenesis (New England Biolabs). This strategy preserved the native signal peptide sequence. The EC1-5/EC3-5 dimer architecture was necessary to eliminate background binding of monomers when EC3-5 was exposed. Cleavage of the signal peptide was confirmed by transfecting and expressing individual constructs and ensuring they were secreted from transfected Expi293 cells (ThermoFisher) and by glycosylation-corrected molecular weight determination using SEC-MALS (Wyatt Technology).

The coding sequences of antibody Fc heterodimerization domains were amplified from the pFUSE-hIgG1-Fc1 plasmid (Invivogen) containing two charged mutations on each opposing domain^57,58^. The SNAPtag sequence was amplified from the pSNAPf vector (NEB). Each of these fragments were cloned into a pFUSE vector (Invivogen) using NEBuilder^®^ HiFi DNA Assembly Master Mix (NEB) along with a tandem 6x polyhistidine tag separated by a GSG linker.

For tip link constructs containing two EC1-5 dimeric binding domains, used in Fig. 3a, the same assembly principle was applied as described above. For full length dimeric constructs, the coding sequences for the entire extracellular domain up to the last hydrophilic amino acid before the transmembrane domain for PCDH15 and CDH23 were used. The PCDH15 R113G mutation was introduced using site-directed mutagenesis (NEB). All constructs were screened for sequence fidelity by Sanger sequencing (GENEWIZ).

Plasmids encoding each half of the final dimerized proteins were co-transfected into HEK Expi293 suspension culture cells in Expi293 medium at a ratio of 4:1 PEI:DNA (w/w), diluted in OptiMem (Invitrogen). A typical transfection used 15 μg of each plasmid with 120 μg PEI in 30 mL of cells at a confluency of 4 x 10^5^ cells/mL. Cells were grown in 8% CO_2_, shaking at 125 RPM. Transfected cells were grown at 37 C for 16-20 hours, then 7 mL Expi293 medium containing 2 mM sodium butyrate and 0.5x penicillin/streptomycin was added, and the cells moved to a 30 C incubator. Cell supernatant was harvested 72-96 hours post-transfection. Cells were removed by centrifugation at 7,000 x g for 15 minutes, and the remaining supernatant was 0.22 μm filtered and stored at 4 C until purification.

Proteins were purified using Ni-NTA or TALON metal affinity purification. First, the supernatant was dialyzed against Tip Link Kinetic buffer (TLK buffer: 20 mM Tris, 125 mM NaCl, 2 mM CaCl_2_, pH 7.4) overnight at 4 C in 3.5 kDa MWCO dialysis tubing (Repligen). 1-1.5 mL of Ni^2+^ Sepharose Excel (GE Life Sciences) or TALON resin (Takara) per 30 mL of supernatant was sedimented in a 10 mL purification column, and then washed with 3 resin bed volumes of PBS followed by 0.5 bed volumes of TLK buffer. The filtered supernatant was next passed through the column by gravity at a rate of ~1-2 mL/min. For Ni-NTA purification, the column was then washed with 3 CV TLK buffer with 15-30 mM imidazole. For TALON purification, the resin was resuspended and washed with 15 CV TLK in batch mode 3 times. In each case, the protein was eluted by resuspending the Ni^2+^ Sepharose with 3 mL TLK buffer with 200 mM imidazole and incubating in a tube on a rotator for 1 hour at 4C. This mixture was then filtered through a 0.2 μM filter and concentrated on a 10 kDa (for EC1-5/EC3-5 proteins) or a 100 kDa (for full length proteins) MWCO column (Vivaspin, GE Healthcare). The protein was buffer exchanged 2-3x using either 7 kDa MWCO Zeba Columns for EC1-5/EC3-5 proteins or 40 kDa MWCO Zeba Columns for full length proteins (ThermoFisher).

Proteins were purified by size exclusion chromatography on a Superose 6 Increase 10/300 column. Purity and molecular weight were confirmed by staining with a SNAP-tag-reactive benzylguanine-Alexa 647 dye (NEB) and running the proteins on a 4-20% NuPAGE Bis-Tris gel. The gels were stained with Krypton protein stain (ThermoFisher) and imaged using a laser gel scanner (GE Typhoon) at 532 nm and 635 nm excitation.

### DNA tethers

Circular ssDNA from the M13 bacteriophage (New England Biolabs) was linearized at a single site using the restriction enzyme BtscI (New England Biolabs) and a site-specific oligonucleotide^59^. A forward primer containing dual 5’ biotins (Integrated DNA Technologies) was purchased and a reverse 5’ benzylguanine primer was synthesized from a primer with a 5’ primary amine on a 12-carbon linker (IDT). Synthesis was accomplished with excess BG-GLA-NHS (NEB) in PBS pH 7.4 for 1 hr at room temperature and buffer exchanged on a Zeba 7k MWCO desalting column. The forward and reverse primers were used to amplify a 2385 base pair DNA fragment from the linearized M13 template using Q5 DNA Polymerase 2x HotStart Master Mix (NEB). The amplified DNA tether was purified using Ampure XP beads at a ratio of 0.7:1 suspended beads to PCR mix and eluted in TLK buffer. The DNA tether was stored at −20 C in aliquots until use.

### DNA-protein tethering complex

Purified SNAP-tagged tip-link fusion proteins (>1 μM) were reacted with >500 nM purified DNA tethers at room temperature for 1.5 hr. Coupling efficiency was assessed by running the DNA-protein complexes on an agarose gel and staining for DNA (Extended Data Fig. 1b). The DNA-protein complex was desalted on Zeba Spin Desalting Columns (ThermoFisher) to remove excess imidazole (if present), and was then purified by Ni-NTA purification using Ni-Sepharose High Performance resin (GE Healthcare) to remove free DNA tethers. The DNA-protein complex was purified as for protein purification, but the washing steps were carried out in the absence of imidazole and was performed entirely in batch mode. The resulting complex was concentrated to 20 μL on a 10-30 kDa MWCO column (VivaSpin, GE Healthcare) and buffer exchanged using Zeba Spin Desalting Columns (ThermoFisher).

### Bead Passivation and Functionalization

Silica microspheres (200 μL of 1% w/v, 3 μM; Bangs Laboratories) were cleaned and hydroxylated by first washing them in a glass tube in MilliQ water, then cleaning with 1% Hellmanex III detergent (Hellma Analytics) by boiling. Beads were then sonicated in the detergent for 5 minutes, then washed and sonicated consecutively with acetone, 1 M KOH, and MilliQ water, and were then rinsed in anhydrous methanol.

The cleaned silica microspheres were aminosilanized by resuspending them in methanol and 20% v/v glacial acetic acid, then adding 1% v/v APTES (Sigma) in a glass tube and mixed by swirling. The solution was then covered with argon gas and rotated end-over-end for 10 min at room temperature. The beads were then rinsed repeatedly with methanol and MilliQ water to remove free aminosilane.

The microspheres were PEGylated using “cloud point” PEGylation^60^. 10 mg of mPEG-5k-NHS and 10 mg of Biotin-PEG-5k-NHS were dissolved in 86.2 μL of 0.5 M Na2SO4 in separate tubes. These reagents were mixed in varying ratios between 1:1 and 1:640 to a final volume of 86.2 μL, giving a cloudy solution. The varying ratios resulted in different surface biotin densities, used later to tune the density of DNA-protein tether on the surface of the beads. 13 μL of 1-M NaHCO3 was added to the mixed PEG solution until the solution just became clear. The cleaned microspheres were pelleted using centrifugation, the supernatant removed, the PEGylation solution added to the beads, and the beads mixed into the solution by carefully pipetting without forming bubbles. The beads were then mixed on an end-over-end rotator with the top end of the tube at a 90° angle from the axis of rotation for 2 hr. The beads were then washed with TLK buffer with 0.02% Tween-20 via successive centrifugation, bubbled with argon and stored at 4 C.

The surfaces of the beads were functionalized with streptavidin to enable binding of biotinylated DNA tethers. 1 μL of PEGylated beads containing surface biotin were diluted 1:50 with TLK buffer + 0.02% Tween-20 + 0.1 mg/mL Roche Blocking Reagent (TLK-TB buffer), and then 50 μM streptavidin was added and allowed to coat the surface for 10 min at room temperature while rotating. One batch of beads was coated with NHS-Alexa 594 labeled streptavidin to enable identification under the microscope via fluorescence. To remove free streptavidin, the beads were then washed five times by consecutive pelleting at 300 x g for 1 min and washing with 200 μL TLK-TB buffer.

The beads were coated with the DNA-protein tethers by resuspending pelleted beads with 3-5 μL of the concentrated, His-purified, DNA-protein complex. The beads were rotated for 1 hr at room temperature and then washed three times with 200 μL TLK-TB buffer and resuspended in 10 μL of TLK-TB buffer. The coated beads were rotated at 4 C until use.

For full-length tip link constructs under some conditions (10 μM Ca^2+^, R113G mutations), it was necessary to perform experiments without the DNA tether in order increase the rate of data collection. For these experiments, fusion proteins were directly biotinylated with BG-Biotin (NEB), buffer exchanged with a Zeba Desalting Column (ThermoFisher) three times, and directly coated onto streptavidin coated microspheres. Individual tethers were verified using a WLC fit to the force-extension traces and tether frequency was kept below 1:10 tethers : attempts.

### Recording Chamber Preparation

Plain 75×25 mm glass microscope slides (VWR) and 22×22 mm glass coverslips (Gold Seal, ThermoFisher) were cleaned by boiling in 1% Hellmanex III (Hellma Analytics) in MilliQ water. The glass was then sonicated for 30 minutes at 37C. After washing thoroughly in MilliQ water, the slides were dried and stored at room temperature in a sealed and dessicated container.

A recording chamber was prepared by laying thin strips of Kapton Tape (DuPont) onto a pre-cleaned microscope slide, then placing and sealing the glass coverslip on top by applying pressure along the tape lines. This created several channels with a height of ~1 mm. Each channel was flushed with 5 mg/mL Roche Blocking reagent dissolved in PBS and allowed to sit in a humidified chamber for 10-30 minutes at room temperature in order to block the glass surfaces and prevent bead sticking. The channel was then flushed with 10-20 chamber volumes of TLK-TB buffer. Each chamber was used immediately after blocking and washing.

1 μL each of the coated beads containing either PCDH15 or CDH23 protein-DNA tethers was diluted 1:100 in TLK-TB and injected into the recording chamber. The chamber was then hermetically sealed with vacuum grease and locked onto the microscope stage of the instrument.

### Instrument Description

The optical tweezer used in these experiments is functionally similar to that described^61^, with several important differences. First, the micropipette was replaced with a second optical trap. This was achieved by using a polarizing beam splitter to split the 1064 nm laser beam into two polarized components. One beam remained stationary as in the previous setup, and the second beam was driven by a piezo mirror with fine control (Mad City Labs, Madison WI), steered in the back focal plane of the objective.

The coverslip containing the recording chambers was mounted on a M-686 XY Stage with Piezoceramic Linear Motors (Physik Instrumente) which was directly attached to the microscope.

### Single Molecule Experiments

Experiments in 2 mM Ca^2+^ were performed at room temperature in TLK-TB buffer. For experiments in Ca^2+^ concentrations below 2 mM, TLK buffer was made without Ca^2+^ or Tween-20 (20mM Tris, 125mM NaCl, pH 7.40). Residual Ca^2+^ from this solution was removed by passing 100 mL of TLK buffer over 5 g of pH-neutralized Chelex 100 resin (BioRad) in a column three times at a flow rate of ~3 mL/min. Ca^2+^ was added back to the Ca^2+^-free TLK buffer to the desired concentration from an analytical stock of 1 M CaCl_2_ solution (Sigma).

Beads were identified on the optical tweezer microscope by Alexa 594 fluorescence, and picked up in each of the trapping beams. Custom software (LabVIEW) was used to control data acquisition and piezo motion. Bead positions were determined in real time at 1400 frames per second with sub-pixel fitting of a polynomial to the dark edges of the beads. The stiffness of each trap was calibrated at 50 mW of laser power using a blur-corrected power spectrum fit^62^. Experiments were performed at 950 mW of laser power, which corresponded to a trap stiffness of ~0.1 pN/nm.

Experiments were performed by bringing the piezo-mirror-driven bead close to the stationary trapped bead for 0.05 – 5 seconds and then retracting at a constant velocity. Formation of an individual tether was cursorily checked by tether length and by displacement of the stationary bead from the center of the trapping beam (Extended Data Fig. 3). When the software detected a single tether, the bead was brought back to the original bead, just close enough that the tether was slack but that there was no chance of forming a second tether. The tether was then quickly pulled out at a constant velocity until the tether ruptured completely. Bead retraction was performed at force loading rates between 0.39 and 377.8 pN s^−1^.

### Data Analysis

Tethering and unbinding events were analyzed individually using custom software (MATLAB). For each set of beads, the average diameter of the two beads was calculated using the polynomial fits to the edges of the beads obtained from the imaging camera. For bead-diameter determination, each set of beads was held stationary in the optical traps for 1400 frames and the diameter of each bead in pixels was determined using the averaged distances from polynomial fits. Pixel distances were converted to nm using a conversion factor of 30.3162 nm/pixel at the defined magnification and verified using a reticule. Extension was measured as the edge-to-edge distance between the two beads in each frame. Individual unbinding traces were fitted with a worm-like chain model to obtain unbinding force, contour length, and persistence length. The distributions of contour lengths and persistence were verified (Extended Data Fig. 5) to ensure only single-molecule bonds were analyzed. Force loading rates were calculated from the linear force regime of each unbinding trace, and the average loading rate at each force was calculated (Extended Data Fig. 4).

Unbinding forces from multiple experiments from at least three individual sets of beads were averaged together for each loading rate tested. The most-probable unbinding force is the force where unbinding probability density is at a maximum. For each set of unbinding forces at a given loading rate we used a kernel smoothing density function (ksdensity in MATLAB) using Gaussian kernels to approximate the unbinding probability density as a function of force^63–65^. The error estimate for each most probable unbinding force was taken to be the kernel bandwidth which was systematically chosen based on the spread of the unbinding force data and the number of rupture forces measured.

Single-bond unbinding data was analyzed using an Evans-Ritchie model for the dynamic strength of molecular adhesion bonds ^22^. Double-bond unbinding data was fit with a multi-state avidity force-dependent avidity model in MATLAB (Supplemental Text).

### Biolayer Interferometry

Proteins were produced and purified in the same manner as for force spectroscopy experiments. Protein molarity was verified using two complementary methods: UV absorbance at 280 in solution, and fluorescence intensity in a SDS-PAGE gel stained with the quantitative Krypton (ThermoFisher) protein stain relative to a BSA standard (ThermoFisher). To create passivated two-dimensional streptavidin-coated biosensors, we used Aminopropylsilane Dip and Read Biosensors (ForteBio), which contain a two-dimensional surface of primary amines. Dry sensors were first hydrated in a variant cloud-point buffer (100 mM HEPES, 0.6M K_2_SO_4_, pH 7.75) in the sensor tray for at least 5 min. All experiments were performed in a 384-well tilted-bottom microplate (Pall ForteBio). Amine-reactive NHS-PEG_12_-Biotin (ThermoFisher) was dissolved to 4 mM in cloud-point buffer and distilled water was added in ~5 μL drops until the buffer just became clarified. Sensors were immediately coated in NHS-PEG_12_-Biotin solution for 2300 seconds on an Octet Red384 instrument (ForteBio) in real time to assess sensor coating efficiency. Sensors whose signals did not group together were discarded. Sensors were then repeatedly and sequentially washed in cloud-point buffer and ultrapure water, then equilibrated in TLK-Tween buffer. Biotinylated sensors were then coated for 200 seconds with 5 μM streptavidin (ProZyme), and washed twice in TLK-Tween buffer. Drift sensors were quenched with 10 μM biocytin (Sigma Aldrich).

In each experiment, CDH23 was biotinylated using a custom-synthesized benzylguanine-PEG15-biotin linker, excess linker was removed using a desalting column or polyhistidine purification, and the proteins were buffer exchanged into TLK-Tween buffer. CDH23 was immobilized to a binding signal of 0.175 - 0.25 nm above baseline in TLK-Tween buffer and washed in at least four separate wells of TLK-Tween buffer before association with PCDH15. Biocytin-quenched drift sensors were dipped into the same concentration of CDH23 for the same length of time as the experimental sensors, in parallel. To assess kinetics, the CDH23+ and CDH23-sensors were dipped into a baseline solution of TLK-Tween buffer, then dipped into a solution of PCDH15 or PCDH15 lacking EC1-2 at 10-300 nM in TLK-Tween buffer to measure association, and finally into the same baseline solution to measure dissociation. PCDH15 and PCDH15 lacking EC1-2 proteins were kept equimolar during each experiment. Experiments were performed using both biological and experimental replicates. Solutions for low Ca^2+^ experiments were prepared as described for single molecule experiments.

PCDH15 proteins lacking EC1-2 were produced by deleted EC1-2 from full-length DNA constructs using the same deletion strategy employed for the EC1-5/EC3-5 dimers used for force spectroscopy. The change in apparent molecular weight was confirmed on an SDS-PAGE gel and via size-exclusion chromatography (Extended Data Fig. 6a).

Each experimental trace was first analyzed using the ForteBio Octet Data Analysis Software 10.0 (Pall ForteBio). Each sensor trace from the biocytin-quenched drift sensor was subtracted from the corresponding experimental sensor, and the subtracted traces were processed using Savitzky-Golay filtering. The drift-corrected sensor trace from the control PCDH15 lacking EC1-2 was then subtracted from the drift-corrected PCDH15 experimental sensor trace. The final subtracted traces were exported into Prism 8.1 software (GraphPad) and each association and dissociation phase were fitted with using single-exponential kinetic equations.

## Acknowledgments

We thank Dr. Marcos Sotomayor for helpful discussion and early conceptualization, and members of the Corey and Wong Laboratories for helpful discussions and for critical reading of the manuscript.

## Author contributions

Conceptualization, E.M.M., A.W., M.A.K., D.P.C., and W.P.W.; Formal Analysis, E.M.M., A.W., W.P.W., and D. P.C.; Funding Acquisition, D.P.C., and W.P.W.; Investigation, E.M.M., A.W., and M.A.K.; Methodology, E.M.M., A.W., D.Y., D.P.C, and W.P.W.; Software, A.W.; Supervision, D.P.C, and W.P.W.; Visualization, E.M.M., A.W., D.P.C, and W.P.W.; Writing – original draft, E.M.M., A.W., D.P.C., and W.P.W.

## Funding

This work was supported by the NIH (F31 DC016199 to E.M.M., R01 DC002281 to D.P.C. and W.P.W., and R35 GM119537 to W.P.W). E.M.M. was a Harvard Medical School Department of Neurobiology Graduate Fellow.

## Competing interests

Authors declare no competing interests.

## Data and materials availability

Methods are as described in the text. Data are available as supplementary materials.

## Materials and Correspondence

Request for materials and correspondence to David P. Corey and Wesley P. Wong.

## Extended Data Figures and Tables

**Extended Data Fig. 1.**
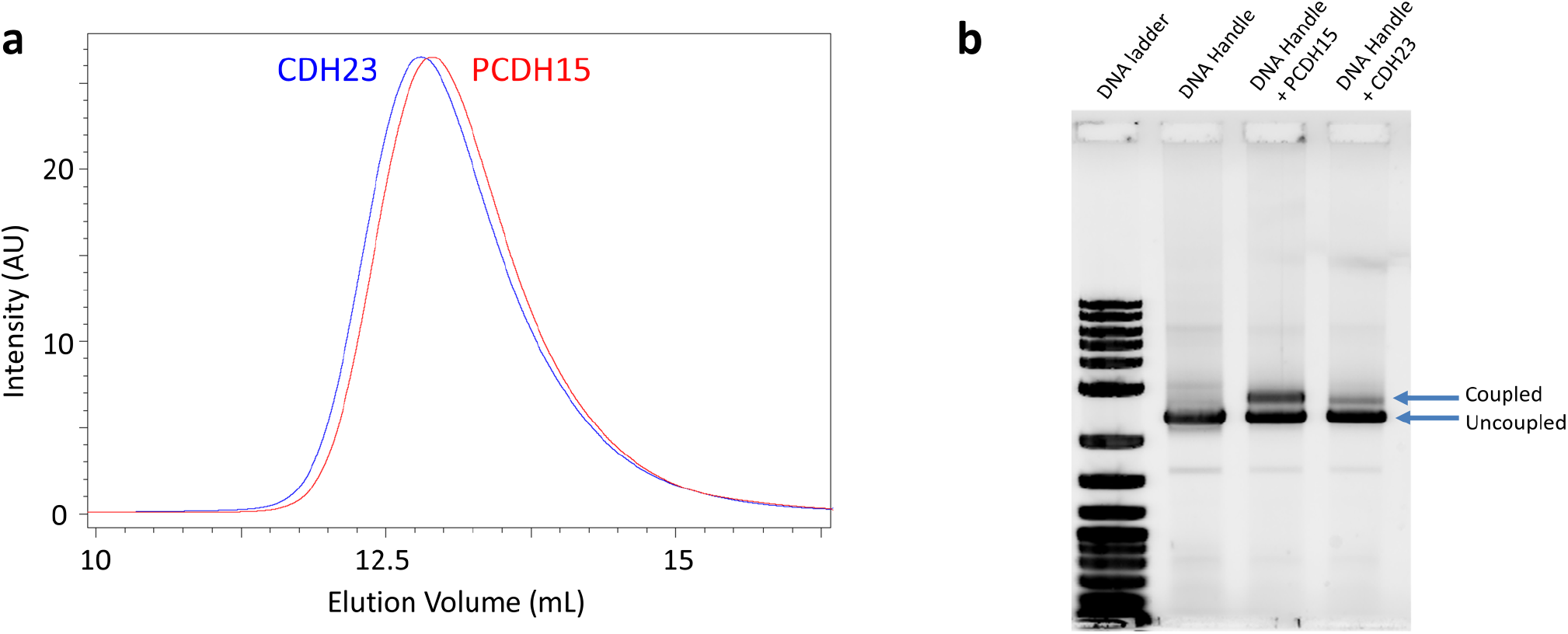
Purification and DNA coupling of single-bond tip link fusion proteins. **a**, Size-exclusion chromatography of single-bond tip link fusion proteins shows monodisperse protein. 100 μL of His-tag purified protein was run on a Superose 6 Increase column at 0.5 mL/min. The fluorescence detector was set to λ_EX_ = 280 nm, λ_EM_ = 340 nm to detect tryptophan fluorescence. **b**, Coupling of purified single-bond fusion proteins to double-stranded DNA handles. DNA handles containing alternative 5’ dual-biotin and 5’ benzylguanine were reacted with SNAP-tagged fusion proteins, and 100 ng of DNA or DNA-protein was run on a 1% agarose gel. The gel was stained with Sybr Gold DNA stain and imaged with a fluorescent gel scanner. DNA-protein complexes are revealed as an apparent shift in molecular weight.

**Extended Data Fig. 2.**
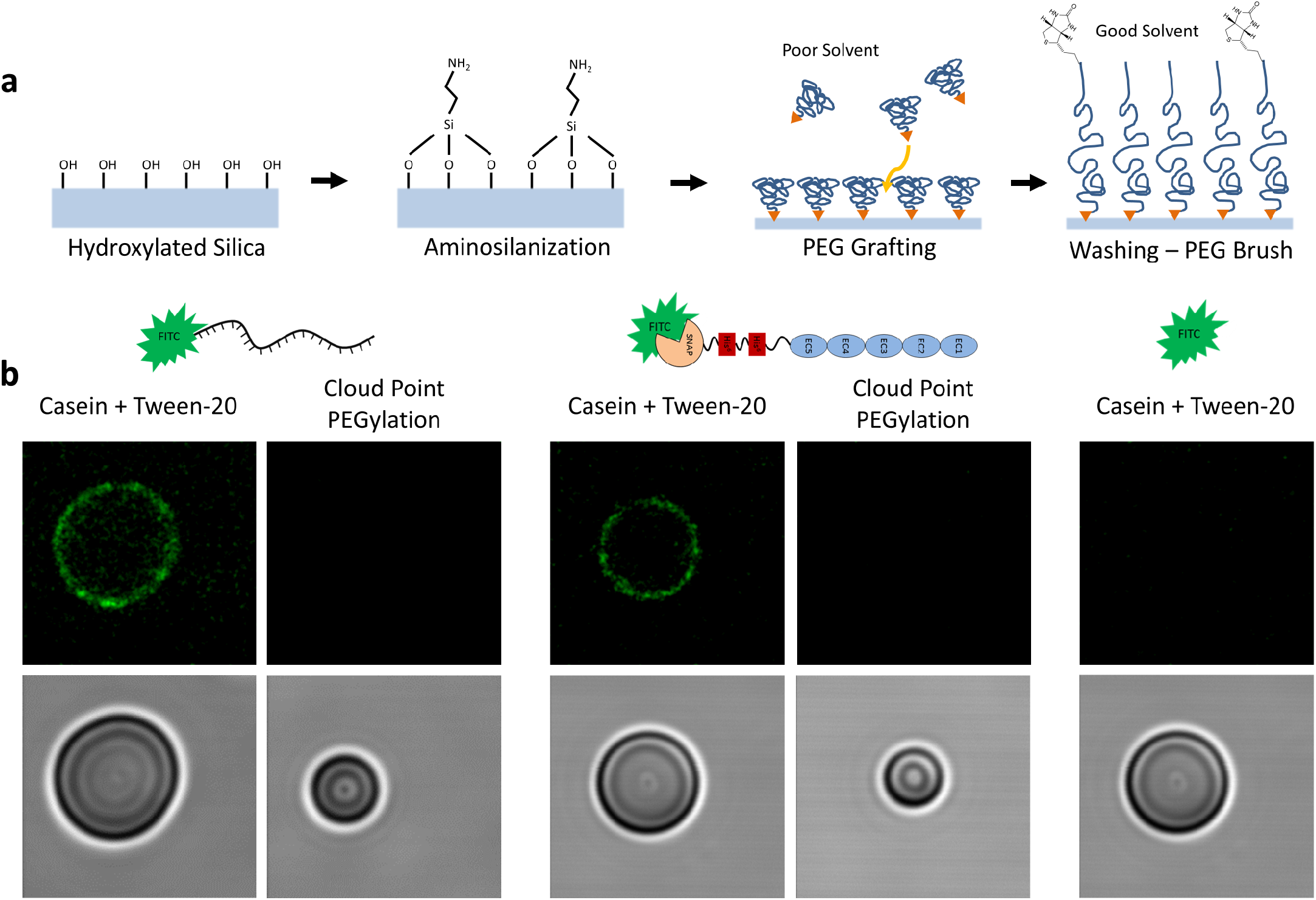
Cloud-point PEG brush grafting for highly passivated surfaces. **a**, Schematic of the chemical grafting process. The silica surface of a microsphere was hydroxylated with a high pH solution and aminosilanized with aminopropylsilane. Amine-reactive NHS-PEG5k-Biotin and NHS-PEG5k-Me was grafted onto the surface in a poor solvent so that the hydrodynamic radius of the PEG molecule was as small as possible. Upon washing the surface with a low-salt saline and detergent solution, the PEG molecules became completely solubilized and formed a PEG brush. **b**, Passivation efficacy assessed using fluorescence microscopy. Single stranded DNA and PCDH15 EC1-5 His SNAP fusion proteins were labeled with a single FITC fluorophore and allowed to interact with either plain silica microspheres in a non-covalent blocking solution of casein and Tween-20 or with cloud-point PEGylated silica microspheres. After washing, the microspheres were imaged using a confocal microscope at an image plane through the center of the microsphere. Cloud-point PEGylation resulted in no discernable non-specific binding.

**Extended Data Fig. 3.**
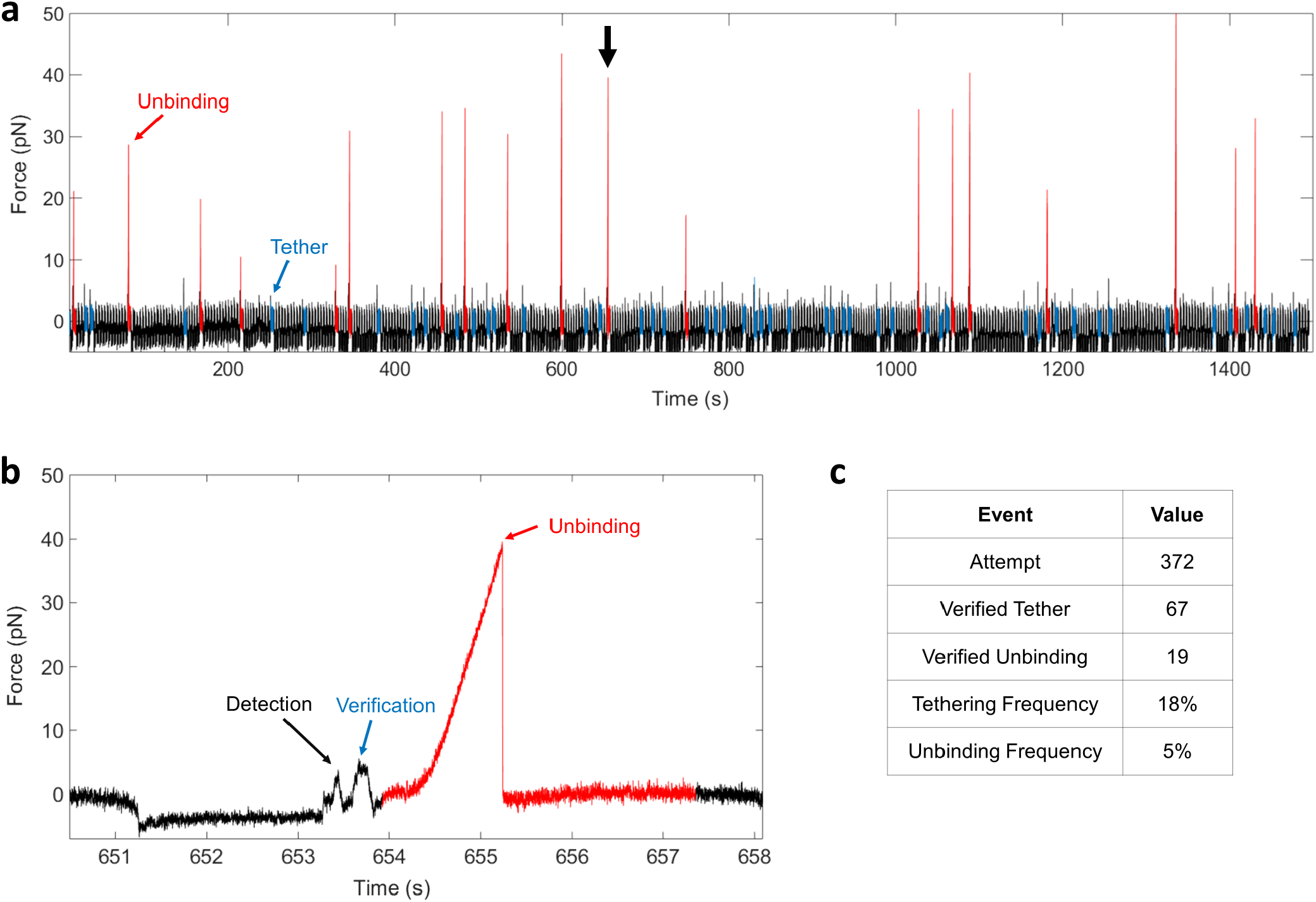
Representative single-molecule rupture events and tethering statistics. **a**, Force-time series of an entire recording session using single-bond proteins (Fig. 1b). Peaks highlighted in red represent single bond unbinding events, and events highlighted in blue represent tethers that were verified by a second pull but unbound before application of the force-loading protocol. **b**, A representative single-molecule unbinding event denoted by the arrow in **a**. Using an automated protocol, beads were brought together for 2 seconds to facilitate bond formation and then pulled apart to detect a tether. Once a tether was detected, the tether length was quickly verified by relaxing and re-applying force, before pulling the tether at a constant velocity of 3000 nm s^−1^ for 3.5 seconds. **c**, Tethering and unbinding statistics for the trace shown in **a**.

**Extended Data Fig. 4.**
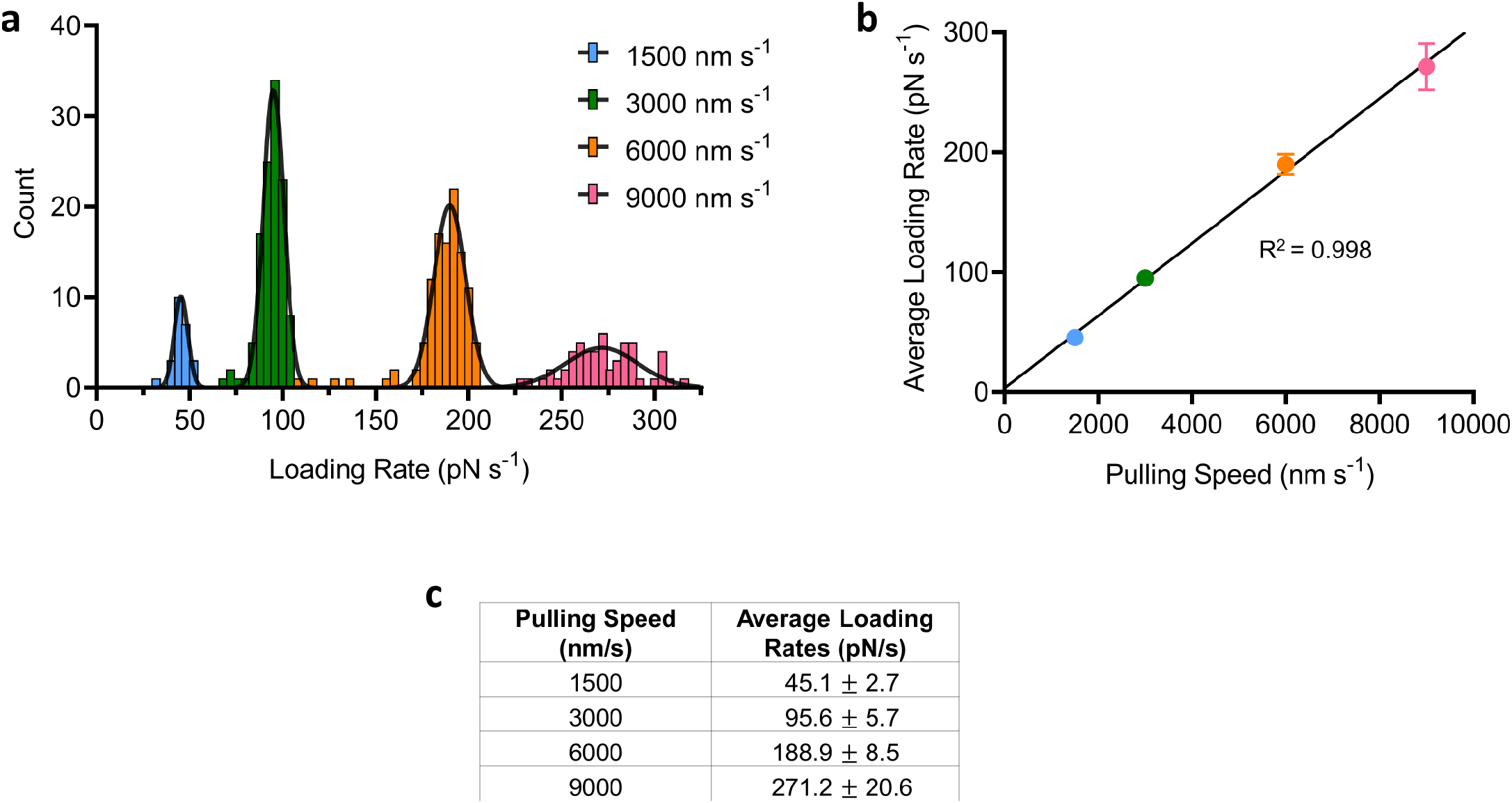
Verification of force loading rates and pulling speeds for single-bond fusion protein unbinding experiments. **a**, Histograms of loading rates obtained from worm-like chain (WLC) fits of single-bond fusion protein unbinding at each of four pulling speeds. Each histogram was fit with a Gaussian function. **b**, Average force loading rate from a plotted against the preset pulling speed. Loading rate increased linearly with pulling speed. Error is the standard deviation from the Gaussian fit. **c**, Data plotted in **b**.

**Extended Data Fig. 5.**
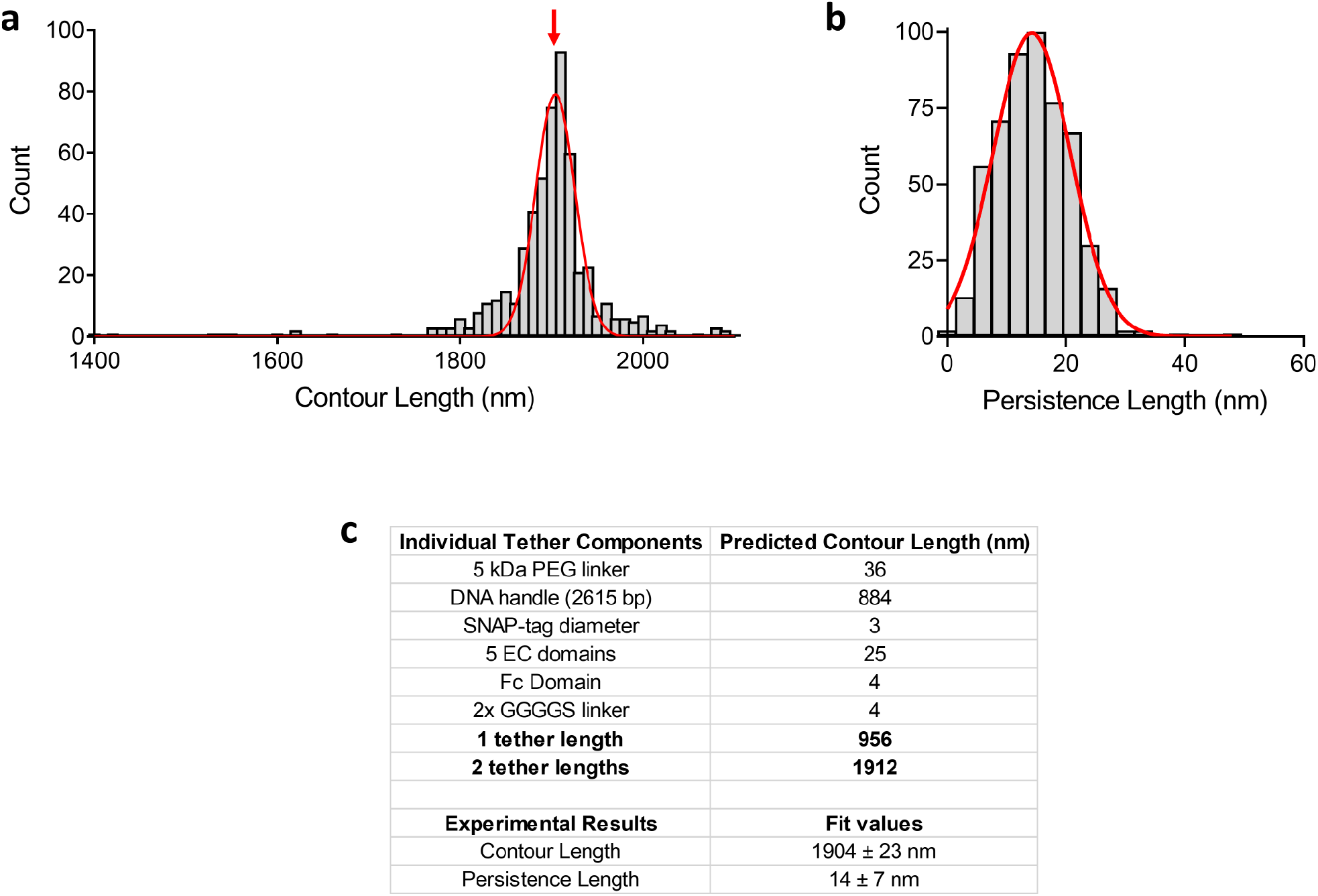
DNA-protein tether characteristics of all single-bond unbinding events. **a**, Histogram of DNA-protein contour lengths obtained from an extensible WLC fit of force-extension data. Events are plotted in 10-nm bins and fit with a Gaussian curve. Contour lengths form a single distribution around the predicted contour length **c**, indicating the measurement of individual tethers containing the correct components. **b**, Histogram of DNA-protein persistence lengths plotted in 3-nm bins and fit with Gaussian curves. The persistence length of a polymer is a metric of its stiffness, and can be used to check for the presence of multiple tethers during an experiment. Two or more tethers measured in parallel should result in discrete, additive persistence lengths. **c**, Calculated theoretical contour length values for each of the individual components in a single tethered complex. 5 kDa PEG has 113 subunits at an average length of 0.318 nm per subunit^66^. DNA subunit length was calculated as 0.338 nm/bp^67^. SNAP-tag diameter was obtained from Protein Data Bank structure PDB 3KZZ. EC domain length was obtained from^8^. GGGGS linker length was calculated from a contour length per amino acid of 0.4 nm^68^.

**Extended Data Fig. 6.**
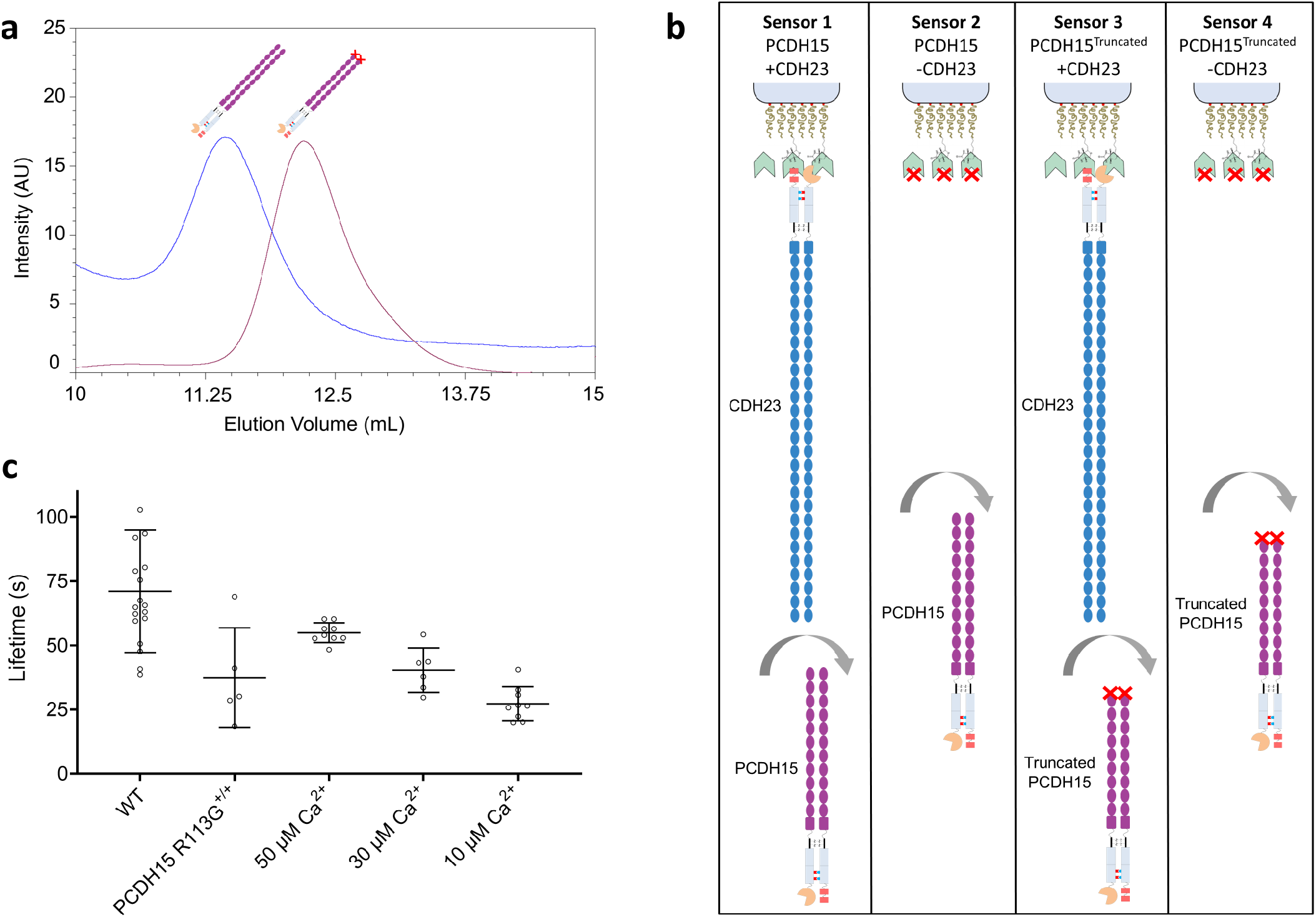
Biolayer interferometry measurements of full-ectodomain tip link proteins. **a**, Size-exclusion chromatography of full-ectodomain, double stranded PCDH15 (blue trace) and the PCDH15 control proteins lacking the EC1-2 binding domains (red trace). Proteins were labeled through their SNAP tags with BG-Alexa 647 and the fluorescence detected with 650 nm excitation and 665 nm emission. Control proteins have a lower apparent molecular weight than WT proteins and each protein forms a single peak. **b**, Biolayer interferometry signal subtraction strategy. Biosensors were coated with PEG-streptavidin and loaded with full-length double-stranded CDH23 (sensors 1–3). The drift-control biosensors were biocytin-quenched and not CDH23-loaded (sensors 2–4). The drift-subtracted truncated control signal (sensor 3 – sensor 4) was then subtracted from drift-subtracted full-length experimental signal (sensor 1 – sensor 2) **c**, Dimeric PCDH15-CDH23 lifetime measurements under different conditions. Dissociation signals from replicate experiments were fit with single exponentials, and the lifetimes measured as the time constants of the fits. The mean and standard deviation are plotted. Both the bond-interface R113G mutation and low Ca^2+^ destabilize the tip-link interaction and decrease its lifetime.

**Extended Data Fig. 7.**
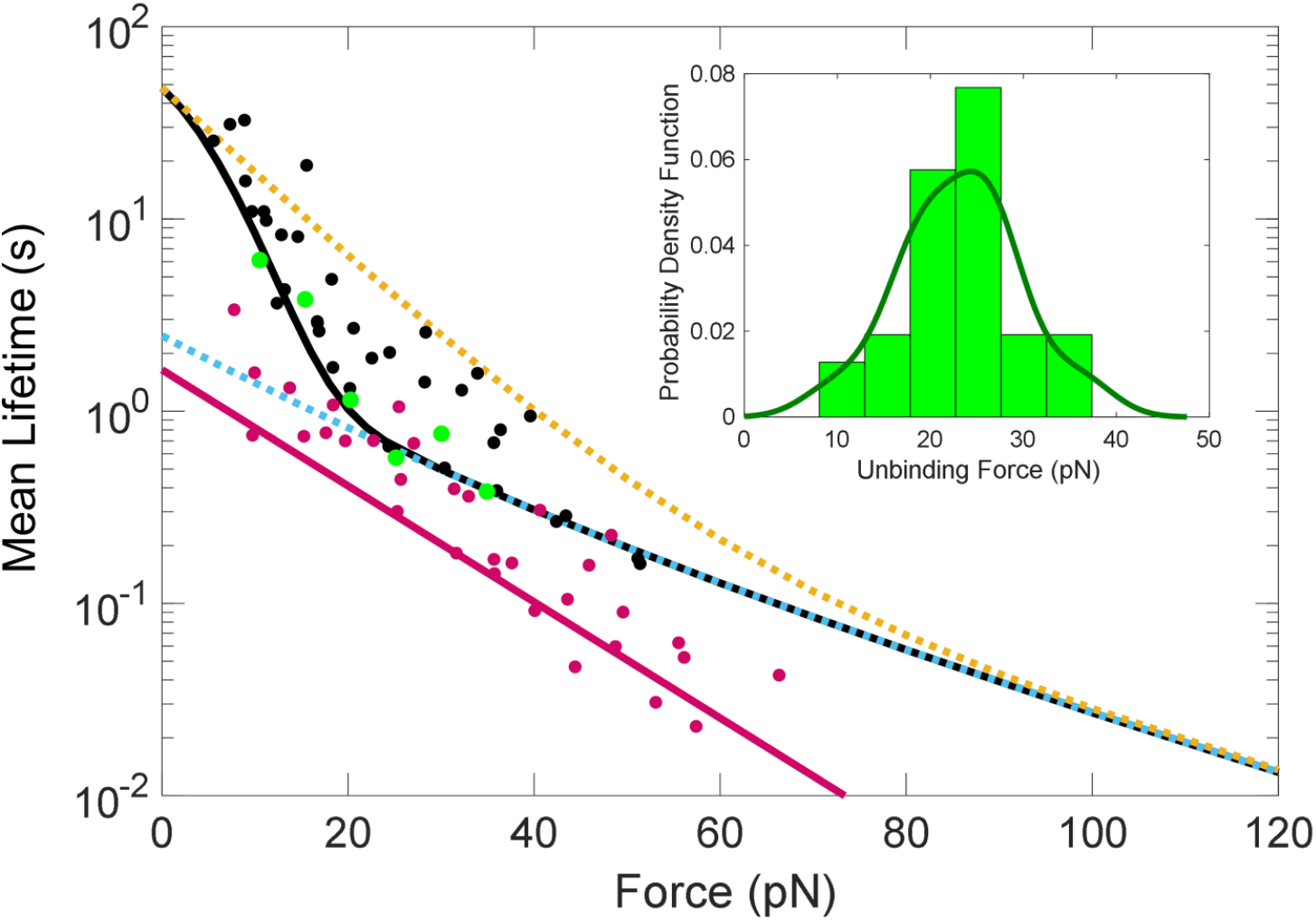
Tip-link lifetimes calculated directly from unbinding force data are consistent with lifetimes calculated from fits by the model. Histograms of single-molecule unbinding forces at each force loading rate were systematically binned using the Freedman–Diaconis rule to yield *N* total bins of width Δ*F* (inset, green bars, 24.1 pN s^−1^, full-length dimer, 2 mM Ca^2+^). A kernel smooth density function is overlaid on the binned unbinding force data. Bin centers were systematically chosen to yield the most likely unbinding force with the maximum number of counts. To estimate the off-rate *k_off_* as a function of force *k_off_* (*F_i_*), we used the following equation^41,69^:

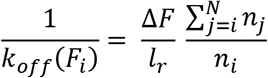 Where *F_i_* is the *i^th^* bin center, *l_r_* is the loading rate, *n_i_* is the number of counts in bin *i*. The estimated mean lifetime as a function of force 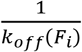 for each bin is plotted on the calculated mean lifetime plot from Fig 2E. These estimates are in close agreement with the mean lifetime predicted from the force-dependent avidity model fit of most-probable unbinding force data. An example of a force histogram is shown as an inset, and calculated lifetimes plotted as green points.

**Extended Data Fig. 8.**
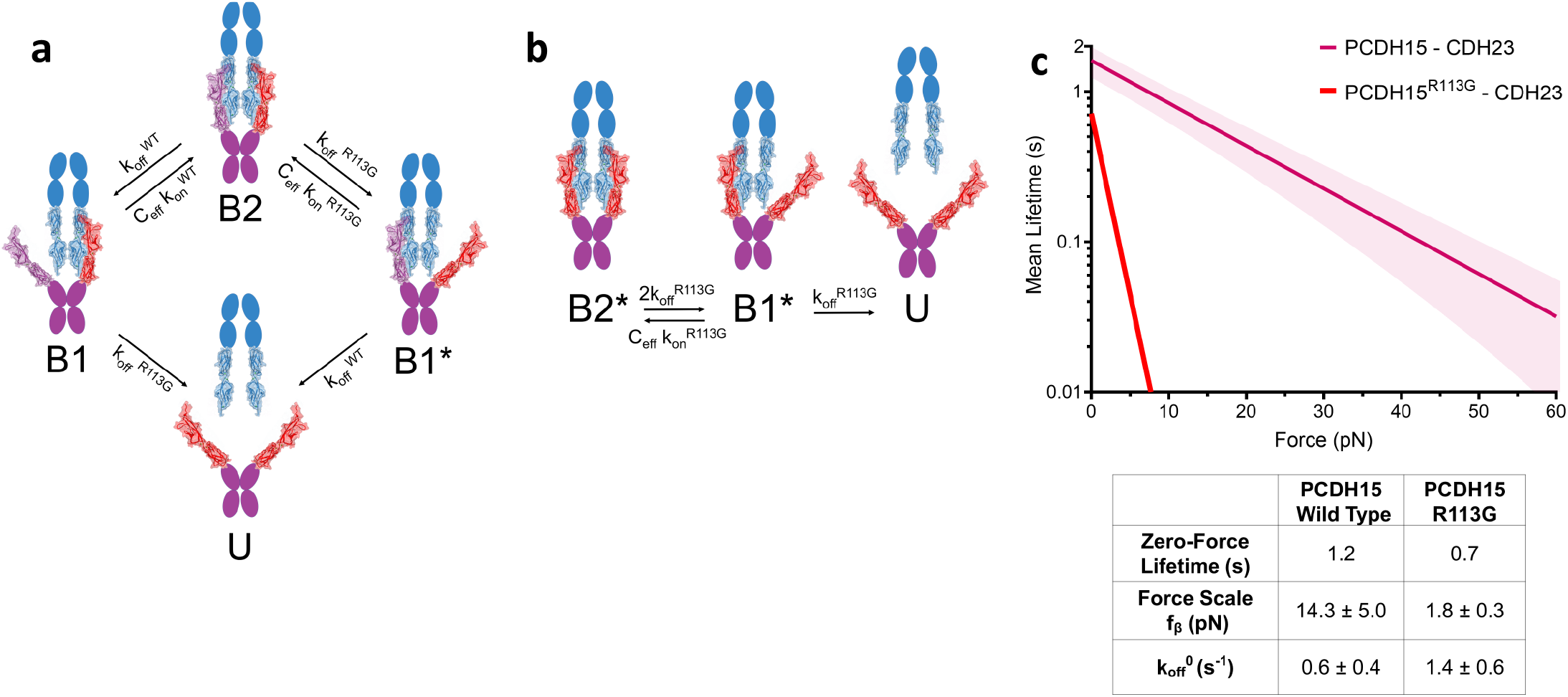
A kinetic model for the PCDH15 R113G human deafness mutation. **a**, A kinetic state diagram representing a tip-link connection heterozygous for the PCDH15 R113G mutation (PCDH15 R113G^+/-^). The doubly-bound complex (B2) can transition to a singly bound state containing a mutated PCDH15 (B1*) or a singly bound state with the wild-type PCDH15 (B1), with rates that differ between mutant and wild type. **b**, If both PCDH15 monomers are mutant, the four-state diagram collapses to the standard three-state diagram, with all-mutant rates. **c**, The single-bond kinetics of the PCDH15 R113G bond, extracted from the best-fit model simultaneously fit to PCDH15 R113G^+/+^ and PCDH15 R113G^+/-^ force spectroscopy data (Fig. 3c), were used to calculate the mean CDH23-PCDH15 single-bond lifetime as a function of force (Supplemental Discussion). The PCDH15 R113G mutation increases both the zero-force off rate (from 0.6 to 1.2 s^−1^) and the force sensitivity of the bond (from 14.3 to 1.8 pN). At a resting tension of 10 pN, a single wild-type bond lasts approximately 1 s, while a single PCDH15 R113G bond lasts less than 10 ms.

**Extended Data Fig. 9.**
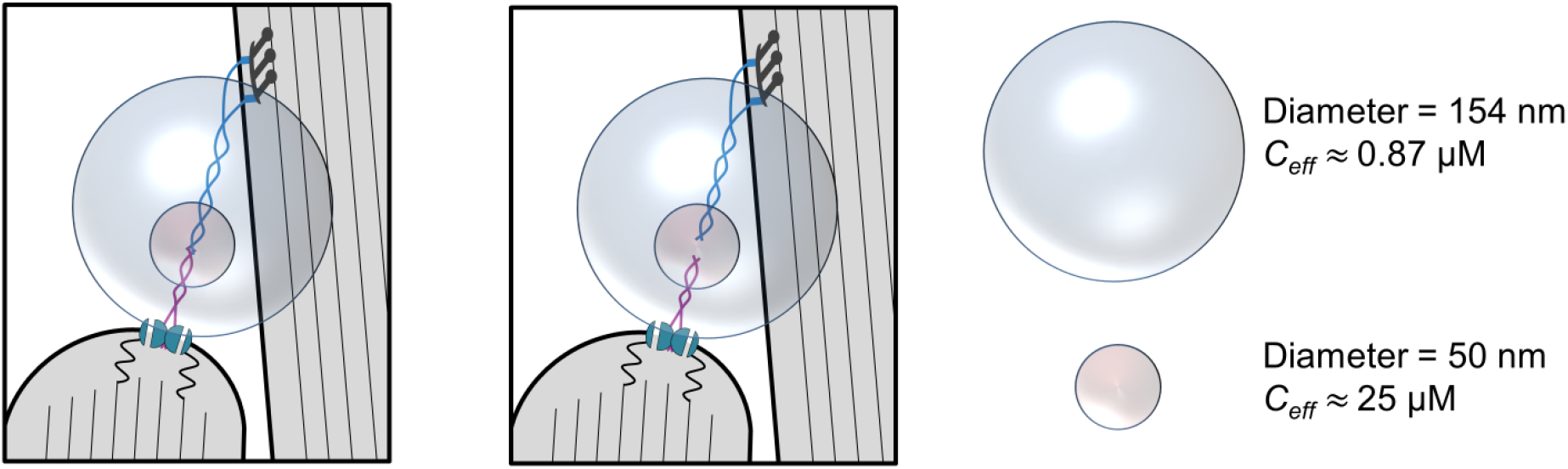
Schematic of the re-formation of the tip link *in vivo*. In order for the re-forming rate to match the measured ~8 second tip-link lifetime, the effective concentration of PCDH15 and CDH23 across the stereocilia gap must be ~0.87 μM (blue sphere, diameter 154 nm). Since the proteins are constrained at their membrane insertion points (Supplementary Discussion), their effective concentration is likely much higher. A more reasonable estimate of the likely space the unbound ends explore is a sphere of 50 nm diameter, corresponding to an effective concentration of 25 μM, and an average reforming time of 0.29 s. If new tip links form near where stereocilia touch, the effective concentration will be still higher.

**Extended Data Table 1.** Kinetic fit parameters of force spectroscopy data.

| Condition | Mean Lifetime (s) | $f_{\beta}$ (pN) | Single-bond $k_{off}$ ($s^{-1}$) | Single-bond $k_{on}$ ($M^{-1} s^{-1}$) $\times 10^4$ | $C_{eff}$ (mM) | $f_c$ (pN) |
| --- | --- | --- | --- | --- | --- | --- |
| EC1-5/3-5 Single Bond<br>2 mM $Ca^{2+}$ | $1.6 \pm 0.4$ | $14.3 \pm 5.0$ | $0.6 \pm 0.4$ | -- | -- | -- |
| EC1-5/3-5 Single Bond<br>50 $\mu M$ $Ca^{2+}$ | $0.9 \pm 0.1$ | $16 \pm 1.5$ | $1.0 \pm 0.2$ | -- | -- | -- |
| Full Length Dimer<br>2 mM $Ca^{2+}$ | $55 \pm 43$ | 14.3 | 0.6 | 7.0 | $0.5 \pm 0.2$ | $10.4 \pm 2.0$ |
| EC1-5/1-5 Dimer<br>2 mM $Ca^{2+}$ | $75 \pm 72$ | 14.3 | 0.6 | 7.0 | $0.7 \pm 0.5$ | $1.9 \pm 0.5$ |
| Full Length Dimer<br>30 $\mu M$ $Ca^{2+}$ | $80 \pm 75$ | $13.8 \pm 4.6$ | $0.4 \pm 0.2$ | 7.0 | $0.4 \pm 0.3$ | $7.8 \pm 0.8$ |
| Full Length Dimer<br>10 $\mu M$ $Ca^{2+}$ | $6 \pm 5$ | $22.0 \pm 15.7$ | $2.6 \pm 0.8$ | 7.0 | $0.9 \pm 0.7$ | $2.9 \pm 1.1$ |
| Full Length Dimer<br>PCDH15 R113G +/- | $14 \pm 6$ | $1.8 \pm 0.3$ | $1.4 \pm 0.6$ | $10 \pm 2$ | 0.5 | $10.5 \pm 3.0$ |
| Full Length Dimer<br>PCDH15 R113G +/+ | $26 \pm 11$ | $1.8 \pm 0.3$ | $1.4 \pm 0.6$ | $10 \pm 2$ | 0.5 | $10.5 \pm 3.0$ |

Values in red are held constant in the fit. R113G +/+ and R113G +/- data were simultaneously fit.

**Extended Data Table 2.**
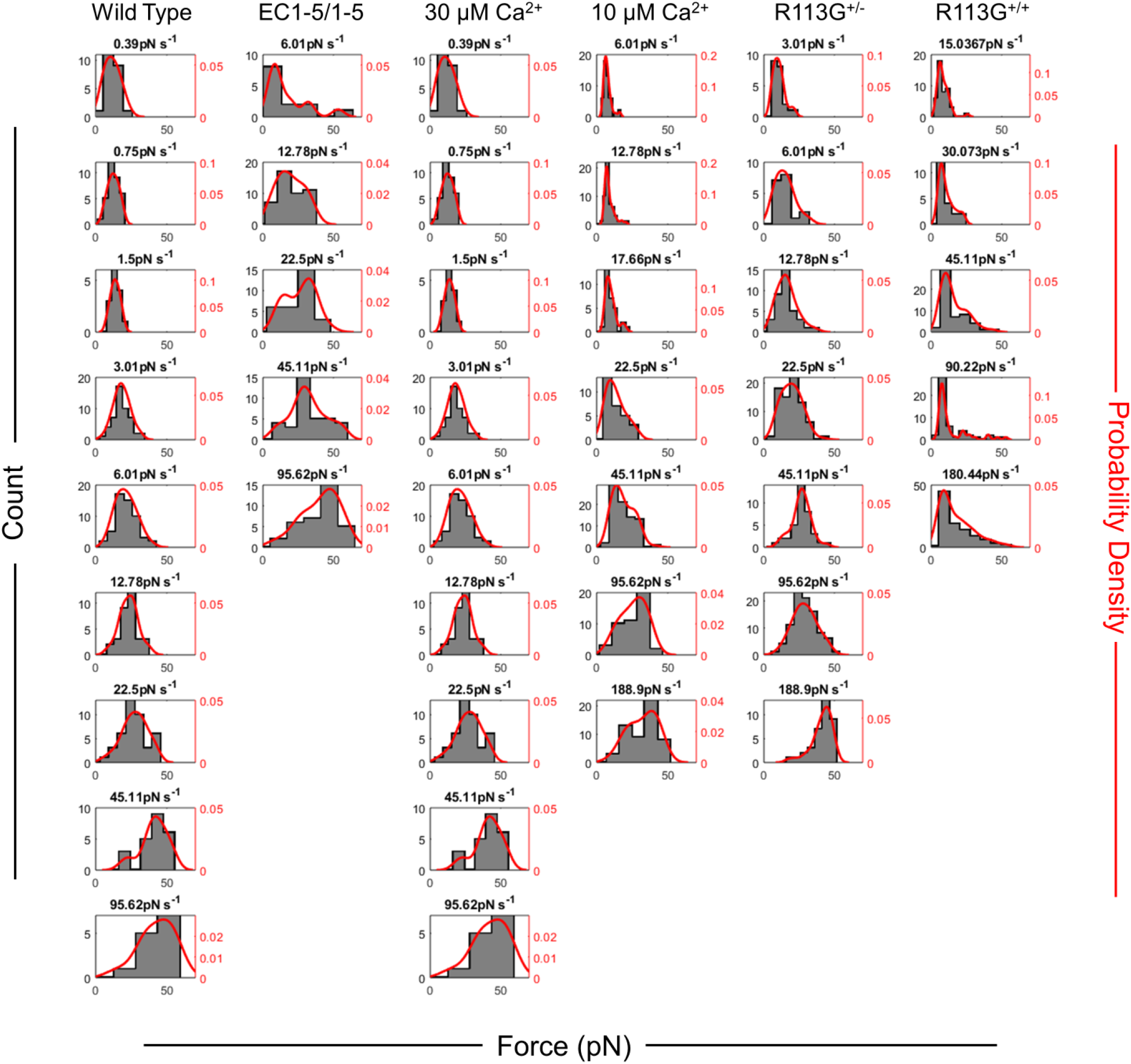
Histograms of all tip-link dimer unbinding data. Histograms of unbinding forces at each force loading rate for each condition were systematically binned using the Freedman-Diaconis rule to yield *N* total bins of width Δ*F*. Bin centers were systematically chosen to yield the most likely unbinding force with the maximum number of counts. A kernel smooth density function is overlaid on the binned unbinding force data. Gaussian kernels were used to approximate the unbinding probability density as a function of force^63-65^.

## Supplementary Information

### Supplemental Discussion

#### A Model for Force-Dependent Avidity

Conforming to the overall structure of the tip-link connection, we modeled the system as two bonds loaded in parallel. In this condition, the force applied to the tip link is distributed equally between the two component bonds. If one bond breaks, the remaining bond assumes the entire force load unless and until the other bond rebinds. We therefore described the two-bond interaction using the following differential equations:

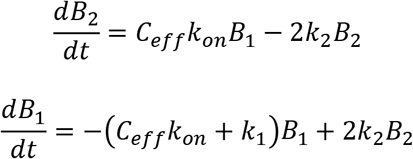

where *B*_2_ represents the fraction of tip links in the double-bound state, and *B*_1_ represents the fraction of tip links in the single-bound state. *k_on_* is the solution on-rate of a single PCDH15 - CDH23 bond. *C_eff_* is the effective concentration of the unbound EC1-2 interface domains. *k*_2_ is the force-dependent off-rate of the PCDH15 - CDH23 bond when there are two bonds splitting the force load (B2➔B1):

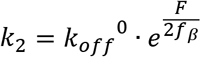

where *k_off_*^0^ is the solution off-rate of a single PCDH15 - CDH23 bond, *F* is the applied force, and *f_β_* is the force which accelerates the off-rate by e-fold.

*k*_1_ is the force dependent off-rate of the PCDH15 - CDH23 bond when there is one bond bearing the entire force (B1➔U):

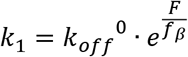

In the case where F=0, these equations can be solved analytically. From this we calculate the mean lifetime *τ* to be:

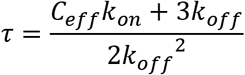

In these experiments, the force changes linearly in time with a loading rate *l_r_*:

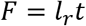

We assume the effective concentration can change as force is applied due to the extensibility of the filaments. Each filament can be thought of as a series of springs each with a spring constant *κ*. If one strand breaks, the free ends separate and tension on the bound strand is doubled, leading to a lower effective concentration *C_eff_*.

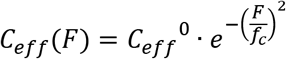

*C_eff_*^0^ is the local effective concentration at zero force; the reduction in *C_eff_* with force is characterized by a force 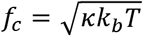 ^41^.

The most probable unbinding force of the dimeric interaction is calculated by finding the most probable unbinding time and multiplying it by the loading rate. The most probable unbinding time is calculated by finding the maximum of the unbinding probability density function:

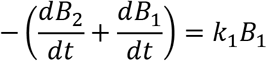

The heterogeneous system is described by the following reaction equations:

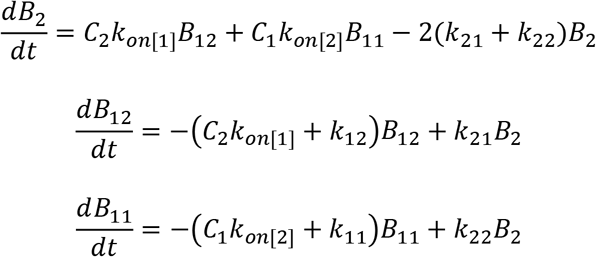

where *B*_2_ represents the fraction of tip links with two bonds. *B*_1*j*_ represents the fraction of tip links with one bond where the index *j* indicates which bond type. *k*_*on*[*j*]_ is the solution on-rate of a single PCDH15 - CDH23 bond. *k_ij_* is the force dependent off-rates of the PCDH15 - CDH23 bond, where the index *i* denotes the number of bonds and *j* denotes the bond type:

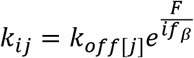

*k*_*off*[*j*]_ is the solution dependent off rate for the *j^th^* bond type. *C_j_* is the force dependent concentration for the *j^th^* bond type.

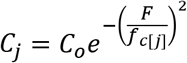

Here we assume that the effective concentration for the *j^th^* bond type decays as force is applied with a characteristic for *f*_*c*[*j*]_.

The most probable unbinding time of the dimeric heterogeneous interaction is calculated by finding the maximum of the unbinding probability density function:

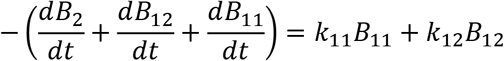

The most probable unbinding time is multiplied by the loading rate to yield the most probable unbinding force. Each model was fit to the data using fminsearch in MATLAB to find the minimum of the *χ^2^*.

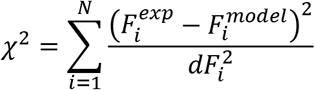

where 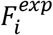 is the *i^th^* experimental most probable unbinding force and 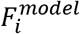 is the most probable unbinding force predicted by the model. 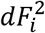 is the estimated square error for the *i^th^* unbinding force. To estimate the error for each fit parameter obtained, we calculated the local curvature of the *χ*^2^ as a function of that fit parameter. The variance (*σ_xi_*)^2^ of the fit parameter *x_i_* is estimated by:

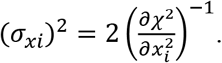

#### Lifetime of the Tip Link in Response to Physiologically Relevant Force Stimuli

Our force spectroscopy experiments were performed with a fixed loading rate, and used to calculate the mean lifetime as a function of static force. However, *in vivo*, auditory stimuli in vertebrate cochleae oscillate hair bundles with a sinusoidal force waveform^51^. Our dynamic-force measurements allow us to make predictions about the behavior of the tip-link connection in oscillating hair bundles. To estimate the tip-link lifetime in response to oscillatory strain, we ran Monte Carlo simulations using the force-dependent off-rates and concentration-dependent on-rates obtained from force spectroscopy (Fig. 4).

What forces are physiologically relevant? In the absence of stimulation, individual tip links are subjected to a resting tension of perhaps 10 pN^35^, established by intracellular myosin motors connected to CDH23. Resting tension maintains transduction channels in their optimally responsive state^70,71^, with a resting open probability P_open_ of ~10%. P_open_ as a function of bundle deflection is well described in hair cells^5,72^, but deducing the range of forces that act on a single transduction complex is more difficult. Estimates for the force operating range of a single mechanotransduction complex, the range of forces that change channel P_open_ from ~10% to ~90%, vary by three-fold or more depending on the organism, type of hair cell, and stimulus method^35,38,54,73–76^. The most sensitive experiments to date suggest that the force operating range is approximately equal to resting tension +5 pN^35,38^, where P_open_ is ~50% at ~12 pN.

We used an established quantitative model of hair bundle mechanics ^53^ to calculate the force on a single tip link in response to an oscillatory bundle deflection, at frequencies from 1 Hz to 10 kHz. We modeled a hair bundle with a gating spring stiffness of 0.6 mN m^−1^ and a geometry factor of *γ* = 0.12 (Fig. 4a-b); thus positive deflection by 133 nm would increase tip-link tension by 9 pN, enough to open essentially all channels if the gating swing is 4 nm^38,54^. We assumed resting tension of 10 pN^35^. In this model, the gating spring does not exert compressive force, so a waveform with high amplitude cannot produce negative tension. For each δt of the simulation (1-10 μs depending on conditions), we calculated the tip-link tension, the force-dependent off-rates, the force-dependent *C_eff_* and the rebinding rate, and then the probability that the tip link remained bound. Each simulation was run until both strands unbound, typically 5-10 s. To obtain a histogram of lifetimes, we ran the Monte Carlo simulation 5000 times for each condition, with phase-randomized sinusoidal stimuli (Fig. 4d-e).

Remarkably, within the range of physiologically-relevant auditory stimulus frequencies (10 Hz – 10 kHz), tip-link lifetime remained largely insensitive both to frequency and amplitude, for peak stereocilia displacements up to 133 nm from rest (~9 pN peak force above resting tension) (Fig. 4d). The insensitivity to frequency over most of the range can be understood if the stimulus period is short compared to the tip-link lifetime (Supplemental Movie 1). Here, there is a higher probability of unbinding during the positive phase, and a higher probability of rebinding during the relaxed negative phase, but overall the lifetime depends on the time spent at each value of tension, and the average tension is no more than resting tension. At larger stimuli the average tension exceeds resting tension and the lifetime is reduced. The reduced lifetimes at low frequency (below 10 Hz) can also be understood: if the period is long compared to lifetime, then an initial positive phase can produce a prolonged high-tension stimulus, and a higher probability that the tip link will rupture within one or a few cycles.

The forces acting on an individual tip link as a function of hair bundle displacement are also modified by the “slow-adaptation” myosin motors that adjust resting tension in tens of milliseconds ^53,54^. To model this, we used a motor climbing rate of 1.6 μm s”^1^ and a motor slip rate of 0.01 s^−1^, yielding a resting tension of 10 pN. When calculating the effect of adaptation, we calculated the slipping and climbing of the adaptation motor in each δt and how that affected tip-link tension.

We found that adaptation enhanced lifetimes both at low frequencies and at high stimulus amplitudes (Fig. 4e). At low frequencies (when the period is longer than the adaptation time constant; here below ~10 Hz), adaptation acts to reduce the stimulus amplitude by reducing tension on the tip link within one stimulus cycle, preventing a high static load and largely negating the effect of a prolonged high tension stimulus (Fig. 4b). At higher frequencies, adaptation reduces resting tension because the slipping rate is faster than the climbing rate, so an oscillatory stimulus causes net relaxation ^70^. With adaptation, even at a peak stimulus deflection of 1000 nm (peak force 72 pN above rest) the lifetime only dropped to 60% of that at resting tension. Overall, these results indicate the tip link is well equipped to transduce auditory stimuli within the force operating range of the transduction channel, for physiological auditory frequencies.

#### Turnover of the Tip Link *In Vivo*

We show the lifetime of an individual tip link at a resting tension of 10 pN is ~8.4 seconds (Fig 2e), suggesting that the tip-link connection is highly dynamic. In order to constantly transduce mechanical stimuli, the tip link must therefore maintain an equilibrium of breaking and re-forming across the gap between stereocilia. Although no measurements to date have been able to discern the kinetic rates of the tip link *in vivo*, several indirect observations have prompted suggestion that the lifetime of a tip link should be much longer – on the order of hours to days. Tip links ruptured by chemical chelation of extracellular Ca^2+^ or exposure to damaging noise stimuli *in vivo* recover with a time constant of approximately 3-6 hours ^11,12^, implying that normal tip-link lifetime should be at least as long as that recovery time.

How can we reconcile a lifetime of 8 s and a recovery time of 6 hr? It may be that the manipulations used to break tip links in recovery experiments – low Ca^2+^ and noise – have additional effects that would inhibit re-forming. First, extracellular Ca^2+^ chelation, typically applied for 15 min, has detrimental effects beyond tip-link rupture, including disassembly of the actin core within shorter rows of stereocilia ^11,12,77^. This may slow the time course of recovery independent of tip-link kinetics. Second, Ca^2+^ chelation and extreme forces lower the energy barrier to tip-link protein unfolding ^39,42^, which may severely damage tip links and associated mechanotransduction components and may require more lengthy refolding before re-forming ^11^. Finally, extended treatment with EGTA or BAPTA abolishes CDH23 immunoreactivity at the tips of bullfrog stereocilia ^36^ and mostly removes CDH23 from the tip-link region in mouse ^12^. It may be that continued inability of a reserve pool to rebind in the presence of 0 Ca^2+^ causes CDH23 to be recycled to the cell body, and then re-establishment of a reserve pool takes ~6 hr.

Our measurements of tip link single-bond kinetics at zero force match well with previously published results ^8,23,32^, and appear to be independent of whether the protein was expressed in bacteria and re-folded (allowing no post-translational modifications) or expressed in mammalian cells. Barring post-translational modifications that occur in hair cells and not in our HEK cells, and that might act to change the affinity of the bond interface, the tip link in hair cells is likely regulated by the same intrinsic single-bond kinetics as that measured *in vitro*. The predominate driving force for increasing a double-stranded tip-link lifetime over that of the single bond should therefore be avidity, the extent of which is determined by the effective concentration of the unbound strands in the single-bound state B1 (Fig. 2a). Could a very high effective concentration, higher than our estimate of ~200 μM at 10 pN, produce a tip-link lifetime of 6 hr? For rebinding to produce a tip-link lifetime of 6 hr, the effective concentration must be 733 mM, and the equivalent volume would correspond to a box 1.3 nm on a side – much smaller than even one EC domain.

Instead, it seems likely that tip-link cadherins can rebind, or new tip links form, on a timescale of seconds. Given a measured on-rate *k_on_* of 1.32 x 10^5^ M^−1^s^−1^ for dimeric PCDH15-CDH23 (Fig. 2b), we can calculate how fast the tip link could re-form. For the re-forming rate to match the ~8 s lifetime, the effective concentration of free CDH23 and PCDH15 must be ~0.87 μM. This concentration roughly corresponds to a sphere with a diameter of 154 nm, about the same as the length of an intact tip link ^7,78,79^ (Extended Data Fig. 9). Since CDH23 and PCDH15 are relatively stiff proteins ^7,40,42^, are complexed with membrane and intracellular proteins ^29,80,81^, and appear to be constrained at their transmembrane domains ^11,12,82^, their N-termini may be constrained to a smaller volume and the effective concentration significantly higher, further increasing the re-forming rate. A more reasonable sphere of 50 nm gives an effective concentration of ~25 μM and a reforming time of just 0.29 s, 29 times faster than the unbinding rate at a resting tension of 10 pN. Moreover, there may well be a reserve pool of CDH23 and PCDH15, near where adjacent stereocilia touch and perhaps already bound, that could be recruited if a recently ruptured tip link cannot re-form ^11^.

### Supplemental Movie 1

A Monte Carlo trajectory of tip-link lifetime using force-dependent off-rates and concentration-dependent on-rates obtained from force spectroscopy of full-length dimers (Fig. 4c). The tip link was deflected with a 10 Hz oscillation. The force as a function of time is plotted, with unbinding events and re-binding events marked with red and green squares respectively. Below is a visualization of the oscillatory strain applied to the filaments. Each filament was visualized as a series of ellipses with an equilibrium spacing of 4 nm and spring constant estimated from the fit of the force spectroscopy data. Movie is presented at 0.18x speed.

